# The effect of mainland dynamics on data and parameter estimates in island biogeography

**DOI:** 10.1101/2022.01.13.476210

**Authors:** Joshua W. Lambert, Pedro Santos Neves, Richèl J.C. Bilderbeek, Luis Valente, Rampal S. Etienne

## Abstract

Understanding macroevolution on islands requires knowledge of the closest relatives of island species on the mainland. The evolutionary relationships between island and mainland species can be reconstructed using phylogenies, to which models can be fitted to understand the dynamical processes of colonisation and diversification. But how much information on the mainland is needed to gain insight into macroevolution on islands? Here we first test whether species turnover on the mainland and incomplete mainland sampling leave recognisable signatures in community phylogenetic data. We find predictable phylogenetic patterns: colonisation times become older and the perceived proportion of endemic species increases as mainland turnover and incomplete knowledge increase. We then analyse the influence of these factors on the inference performance of the island biogeography model DAISIE, a whole-island community phylogenetic model that assumes that mainland species do not diversify, and that the mainland is fully sampled in the phylogeny. We find that colonisation and diversification rate are estimated with little bias in the presence of mainland extinction and incomplete sampling. By contrast, the rate of anagenesis is overestimated under high levels of mainland extinction and incomplete sampling, because these increase the perceived level of island endemism. We conclude that community-wide phylogenetic and endemism datasets of island species carry a signature of mainland extinction and sampling. The robustness of parameter estimates suggests that island diversification and colonisation can be studied even with limited knowledge of mainland dynamics.

## Introduction

Islands have become known as natural laboratories for the study of biodiversity, because of their isolated nature, young geological age, and low ecological complexity relative to the continent (Whittaker et al., 2017). Oceanic islands — those that form *de novo* without any initial biodiversity — are ideal for studying community assembly from first principles as species have to colonise the depauperate island (MacArthur and Wilson, 1967; Mittelbach and Schemske, 2015). Biodiversity on islands is governed by the dynamics of colonisation of the island, extinction, as well as *in situ* speciation (cladogenesis), which are all influenced by island area and isolation (MacArthur and Wilson, 1967; Losos and Schluter, 2000; Kisel and Barraclough, 2010; Valente et al., 2020).

Biodiversity patterns and evolutionary processes on islands are increasingly being investigated using phylogenetic data on island species (Rabosky and Glor, 2010; Hua and Bromham, 2020). However, a key issue that has never been formally evaluated is how main-land species turnover and sampling of mainland species (ancestors or close relatives of island taxa) influence island phylogenetic data and, consequently, our ability to make meaningful and accurate inferences on insular evolutionary processes. First, mainland processes can influence our ability to determine the timing of colonisation of an island (Fig. 1). Island biogeography theory predicts that the divergence of neutral loci between mainland and island populations provides an estimate of the time of colonisation (Johnson et al., 2000); and genetic signatures of island colonisation are found in empirical studies (Sendell-Price et al., 2021). The time of colonisation is then the time the island lineage diverged from its closest (extinct or extant) mainland relative (i.e. species’ phylogenetic stem age), but because accurate data on extinct species on the mainland is rarely available, the time of colonisation is often estimated to be less recent than it actually is (Fig. 1). If the reconstructed phylogeny is densely populated with branching events and the extinction rate is low, the error is expected to be small (Fig. 1A), but if extinction is high or speciation rate is low on the mainland, the approximated colonisation time may be far from the true value as the time back to the previous branching time may be deep in the past (Fig. 1B). In some cases, hereafter termed *maximum island age colonisations*, the divergence time to the closest sampled or extant mainland species in the phylogeny is older than the island, in which case the colonisation time is approximated by the island age as species could evidently not have colonised earlier. A similar effect is produced by incomplete sampling of the mainland species pool (Fig. S1). Incomplete sampling, for example due to incomplete knowledge (e.g., the closest mainland ancestor is unknown) or unavailability of samples, can lead to phylogenies where the stem age of an island lineage gives the impression of an older colonisation time than the actual colonisation time. If there is *in situ* divergence on the island, the crown age can also be used as a time of colonisation (García-Verdugo et al., 2019). Another approach calculates the probability of colonisation between two calibration points, for example the stem and crown age of a radiation, or, in the absence of information on these ages, between the age of the island and the present (Valente et al., 2015, 2019). Finally, colonisation times can also be estimated via a probabilistic ancestral geographic species range reconstruction using a specified model of species range evolution, a phylogeny, and current range data (Ree et al., 2005; Goldberg et al., 2011; Hua and Bromham, 2020). However, this probabilistic method also suffers from a lack of data on extinct species (Gascuel and Steel, 2020).

**Figure 1:**
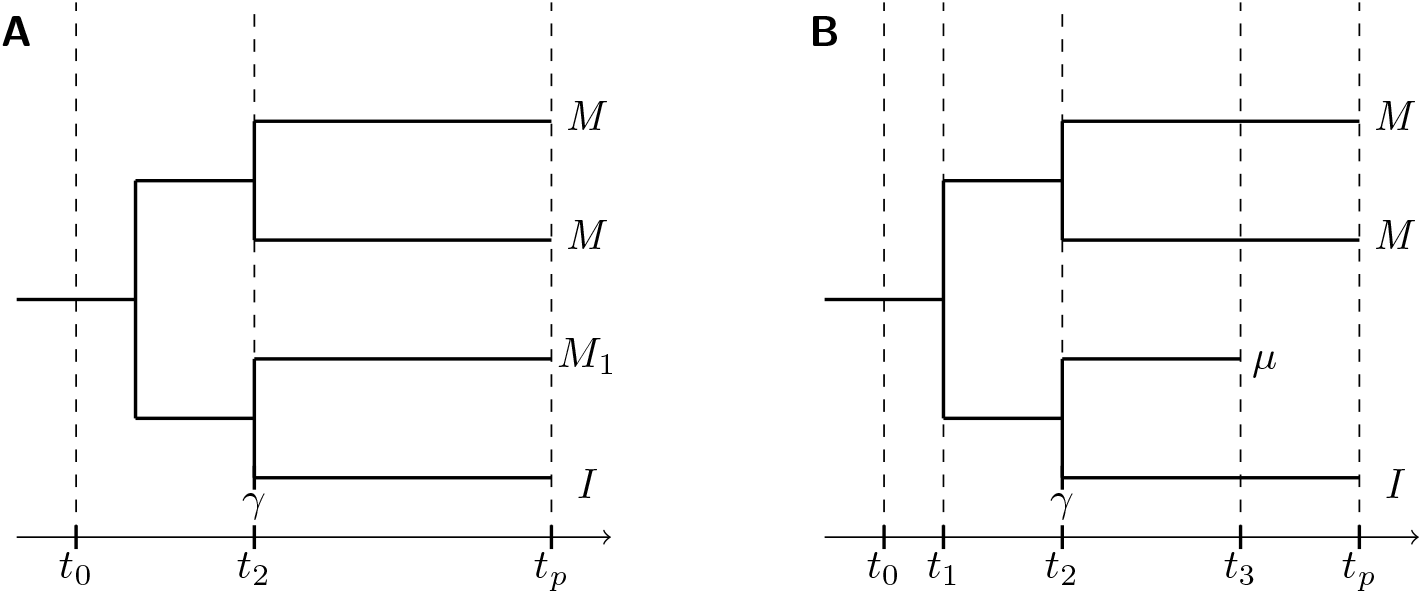
Approximating the colonisation time from the branching time of the island (endemic) species (*I*) with its most closely related mainland species (*M*_1_). The island emerges at *t*_0_, there is a speciation event on the mainland at *t*_1_, a species colonises the island at *t*_2_ (*γ*), leading to a divergence between the island and mainland lineages and causing a branching time at *t*_2_; *t*_*p*_ is the present time. (A) If the mainland sister species is extant, the colonisation time is correctly described by the divergence time. (B) Extinction on the mainland causes overestimation of the colonisation time. The colonisation of a mainland species to the island occurs at *t*_2_, but as the sister species of the colonist goes extinct at *t*_3_ (*μ*), the colonisation time inferred from divergence from the most closely related sister species (*M*) is *t*_1_.

A second key effect of mainland dynamics and incomplete sampling is that these can influence the level of (perceived) endemism on the island. If species go extinct rapidly on the mainland, this can lead to higher proportions of endemic species on islands. Likewise, incomplete knowledge of the mainland can lead to the perception that some species are endemic to the island, when they are in fact also present on the mainland. An increase in the perceived proportion of endemic species may in turn suggest that speciation rates on the island are higher than they really were.

The above-mentioned effects of mainland turnover and incomplete sampling may lead to signatures or errors in the phylogenetic data, which in turn are likely to influence the performance of phylogenetic models. Biodiversity patterns can be studied using the phylogeny of an island clade that radiated after colonisation. Fitting a set of candidate phylogenetic models can then help understand the tempo and mode of diversification. It is commonly observed from phylogenetic data that clades undergo an early-burst radiation with a high rate of speciation, before a slow-down in diversification (Phillimore and Price, 2008; Rabosky and Glor, 2010). This slowdown has been attributed to diversity-dependent diversification, due to, for example niche-filling of species within a clade (Etienne et al., 2012) (but see Etienne and Rosindell, 2012; Moen and Morlon, 2014, for alternative explanations). A multitude of methods have been developed to determine whether diversification has occurred at a constant rate through time, depends on the diversity of the clade through time, or on some environmental or phenotypic factor (Morlon, 2014). However, a limitation with these approaches is that they require radiations of a substantial size (*>* 50 species) in order to have sufficient power for distinguishing the aforementioned phylogenetic models (Etienne et al., 2016). Islands are thought to be prime environments to allow radiations into novel niches with many well-studied radiations (see Losos and Ricklefs, 2009), but island radiations are the exception rather than the rule, being outnumbered by species-poor or single-species clades, limiting the application of single-clade phylogenetic models (Valente et al., 2020).

To circumvent these limitations, one can investigate the colonisation and diversification dynamics of an entire insular community taking into account all species of a particular taxon or ecological guild, irrespective of whether they have diversified on the islands. The Dynamic Assembly of Island biota through Speciation, Immigration and Extinction (DAISIE) model does just that; it is a methodological framework to simulate and make inferences on whole island communities containing multiple colonisation events (Valente et al., 2015). DAISIE uses phylogenetic data of an island community to estimate rates of colonisation, speciation (cladogenesis and anagenesis), and extinction on the island as well as a clade-level carrying capacity (i.e. diversity-dependent colonisation and diversification acts within clades, not between clades). Speciation consists of two processes, firstly, cladogenesis, which defines the rate of *in situ* speciation with lineage splitting on the island, and secondly, anagenesis, which defines the rate at which non-endemic species on the island diverge from their main-land population in the process of becoming endemic. By incorporating information on the age of the island, timing of species colonisation, species endemicity as well as phylogenetic branching times, DAISIE can test whether island communities assemble at a constant rate or are influenced by negative diversity-dependent feedback. Only cladogenesis and colonisation are currently assumed to be diversity-dependent, because diversity-dependent extinction is not easily or commonly detected from molecular phylogenies (Rabosky and Lovette, 2008; Burin et al., 2019), and anagenesis occurs through divergence from a mainland population and thus is less likely to be limited by the diversity of species on the island. Application of DAISIE to several empirical systems has shown that island communities can exhibit both equilibrial and non-equilibrial diversification dynamics simultaneously (Valente et al., 2015); that colonisation and diversification depend on area and isolation (Valente et al., 2020); a shift in colonisation results from a change in island size (Hauffe et al., 2020); and that biodiversity lost through anthropogenic extinction requires several million years to replenish assuming pre-human rates of colonisation and diversification (Valente et al., 2017a, 2019). These results are validated with evidence that phylogenetic data greatly improves estimates of diversification and colonisation (Valente et al., 2018).

Here, we investigate to what extent mainland dynamics and incomplete sampling can influence: 1) phylogenetic datasets of islands species, 2) estimates of rates of colonisation, cladogenesis, extinction, and anagenesis on the island, and 3) estimates of the island carrying capacity. We test how mainland extinction and incomplete sampling (due to not sampling known species as well as not sampling undiscovered/unknown species) affect the performance of the phylogenetic inference model DAISIE in estimating diversity patterns through the processes of the colonisation, speciation and extinction. The bias in colonisation rate is expected to be proportional to the rate of mainland extinction: as more species go extinct, the colonisation time from the phylogenetic tree gets pushed back into the past (Fig. 1B). Mainland extinction also influences the rate of anagenesis, as species may become endemic as a result of the mainland population going extinct (paleoendemism) and not because the species has diverged in allopatry on the island from its mainland population. The rate of cladogenesis is potentially biased downward for the same reason as colonisation: the time a species reached the island is biased towards the past, and thus the same number of *in situ* speciation events occur in a longer period of time. Because cladogenesis is usually estimated under a birth-death process the rate of extinction will also be influenced (Etienne et al., 2012). Additionally, if multiple colonists derive from a now extinct mainland species, it appears from the phylogenetic data that these are descendants of a single colonist, because they are more closely related to each other than any extant mainland species, thus potentially resulting in an artifactual increase in the estimates of cladogenesis, while decreasing colonisation.

## Methods

### Dynamic mainland DAISIE model

In the original DAISIE simulation (and inference) model it is assumed that the mainland species pool (i.e. all species that are potential colonisers of the island) is composed of a constant number of independent species which do not undergo speciation or extinction. A mainland species can only immigrate to the island (Fig. 2A). Here we relax this assumption by extending the DAISIE simulation model to include mainland dynamics. The dynamic mainland model introduced here has the same island processes as DAISIE: colonisation (at rate *γ*) from a mainland with *M* species, cladogenesis (at rate *λ*^*c*^), extinction (at rate *μ*), anagenesis (at rate *λ*^*a*^) and a carrying capacity (*K’*). We simulate the mainland with a species-level Moran process (Moran, 1958), i.e. extinction is immediately followed by a cladogenetic speciation event of a random species on the mainland, and thus the mainland species pool has a fixed size (*M*), as in the original DAISIE model. Thus, we only make the mainland dynamic, but otherwise stay as close to the original DAISIE simulation and inference model as possible. The Moran process is approximately equivalent to the diversity-dependent birth-death process when the diversity is at the carrying capacity (equilibrium) and the birth rate is large. The Moran process may influence diversity-dependence on the island if multiple colonising lineages belong to the same mainland clade (phylogenetic non-independence). The mainland extinction events are independent random variables which have an exponentially distributed waiting time, and thus the number of extinction events that occur across all the mainland species is a Poisson point process (Ross, 1996). While the Moran process with instant replacement may seem unrealistic, it has been shown that some clades diversify in such a coalescence-like process (Morlon et al., 2010). But as stated above, the most important reason to choose the Moran process is to isolate the effect of mainland dynamics rather than the effect of mainland size variability. The static phylogenetically independent mainland pool assumed by the DAISIE inference model is a convenient property required for tractability of the computation of the likelihood. It means that each island colonist diversifies under its own diversity-dependent process (i.e. it and its descendants do not compete with other island colonists and their descendants). However, in the dynamic mainland model when two related (non-independent) species colonise the island they do influence each other’s diversification.

**Figure 2:**
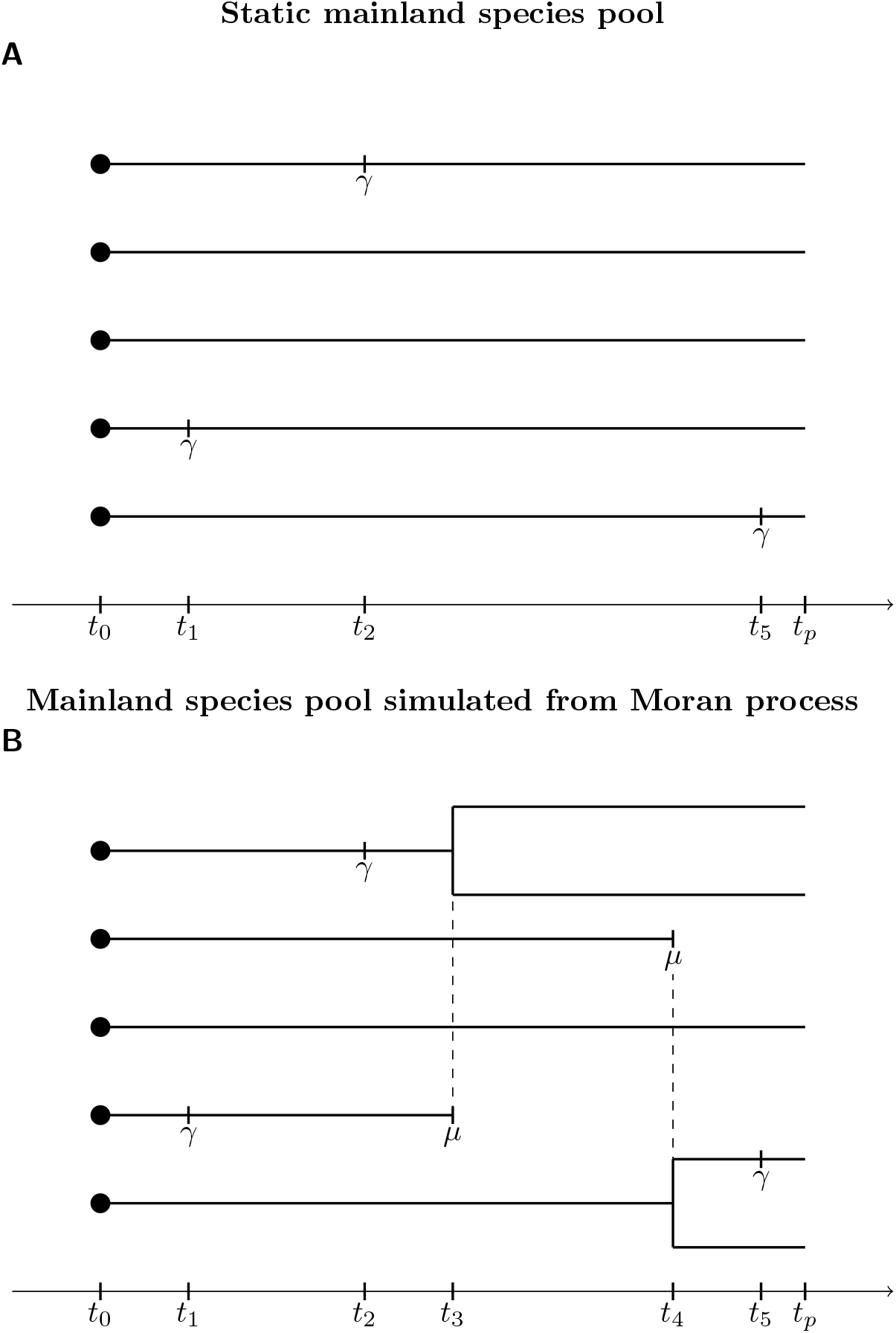
Example of the mainland simulations in the DAISIE model. (A) original DAISIE simulation with static mainland species pool, where species cannot go extinct; species immigrate to the island (*γ* at times *t*_1_, *t*_2_ and *t*_5_), (B) DAISIE simulation of this paper where mainland dynamics are described by a Moran process, where species can go extinct or speciate; species immigrate to the island (*γ* at time *t*_1_, *t*_2_ and *t*_5_). Extinction is denoted by *μ* and connected by a dashed line to the immediately subsequent speciation event shown by a branching event (*t*_3_ and *t*_4_). Species that immigrate as part of the same mainland clade experience the same diversity-dependent process on the island. In both simulations the mainland species remains on the mainland after it immigrates to the island, and can re-immigrate at a later time if it has not gone extinct. After speciation on the mainland, the identity of the species changes, and two new mainland species are formed. Both scenarios have a timeline starting at *t*_0_, with events at *t*_*n*_, and ending at the present *t*_*p*_.

We assume that there are *M* mainland species at the start of the simulation and at any time in the past (zero-sum assumption). They belong to the same broad group as the island community (e.g. terrestrial birds of South America for the terrestrial birds of the Galápagos (Valente et al., 2015)). With mainland extinction, *M* is smaller than the total number of species that could have colonised the island since it arose, because some of the potential colonisers will have gone extinct before the present. Still, the total species-time (i.e. number of species multiplied by time extant) available for colonisation is *M × t*_0_ (where *t*_0_ is the island age). The mainland only needs to be simulated during the time frame in which the island exists and can be colonised, because the maximum age, before the present, at which a species can colonise the island is the age of the island itself (*maximum island age colonisation*). Extinctions on the mainland before the island’s existence do not influence the colonisation times input into the DAISIE inference model (Fig. S2).

### Incomplete sampling and undiscovered species of the mainland pool

Phylogenetic trees are often incomplete, in other words, they are composed of only a sub-sample of all species in a taxonomic group. There are several methods for incorporating unsampled island species when estimating macroevolutionary rates from reconstructed phylogenies using models such as DAISIE and other birth-death models (Nee et al., 1994; Stadler, 2009; Höhna et al., 2011; Etienne et al., 2012; Valente et al., 2015) and previous research has shown the bias of ignoring incomplete sampling from inference on phylogenetic trees (Pybus and Harvey, 2000; Cusimano and Renner, 2010). However, a method for accounting for the missing species on the mainland has not yet been incorporated into an inference model such as DAISIE. Therefore, we assume for this study that all the island species are known and sampled, and focus on the phylogenetic consequences of incomplete sampling on the mainland. Such sampling is analogous to a mass extinction having occurred immediately before the present (Nee et al., 1994), so we expect incomplete sampling to have a similar effect as mainland extinction.

The missing mainland species include those that are taxonomically known but not sampled in the phylogenetic tree, hereafter termed unsampled species, and species that are yet to be discovered, hereafter termed undiscovered species. We inspect the effect of each of these on phylogenetic data. To test the effect of incomplete sampling of the mainland pool, we used a so-called *ρ* sampling approach to prune away tips from the phylogenies on the mainland with a probability *ρ*. Each species on the mainland has an equal probability (*ρ*) of being pruned from the data set under the assumption of random sampling. Undiscovered species have a different effect on the phylogenetic data than unsampled known species. When the mainland ancestor of an island species is unsampled but known to exist, we lose information on the colonisation time, but the island species is correctly classified as non-endemic. When the mainland population of an island species is undiscovered, we also do not have information on the precise colonisation time, but, in addition, a non-endemic island species will be erroneously classified as endemic.

Incomplete sampling of the mainland, in the case of either unknown or undiscovered, is expected to lead to a similar bias as mainland extinction because missing species will cause the colonisation time to be pushed further into the past (Fig. S1). The more incomplete the sampling (lower *ρ*), the more error is expected in the rates.

### Simulating data and parameter estimation bias

To determine the amount of error made given the incomplete information – either via incomplete knowledge on extinct species, or incomplete knowledge on extant species from unsampled known species or undiscovered species, we simulated a complete phylogenetic data set and produced three incomplete data sets by pruning out information from the complete set. The complete information data set is termed *ideal*. This has the exact time of colonisation and correct endemicity status for every lineage. The incomplete information data sets are termed *empirical* as this is the information expected to be available to empiricists. The first *empirical* data set is influenced by mainland extinction (*empirical*_*ME*_). The second and third *empirical* data sets are subject to incomplete phylogenetic sampling of mainland species *empirical*_*US*_ (i.e. *empirical unsampled*), and a failure to discover mainland species *empirical*_*UD*_ (i.e. *empirical undiscovered*), without the influence of mainland extinction. There may be interactions between mainland extinction and incomplete sampling, but here, for simplicity, we examine their effects separately.

We simulated the DAISIE model with a dynamic mainland using the Doob-Gillespie algorithm in continuous time (Gillespie, 1977) and implemented it in the R package DAISIEmainland version 1.0.0 (Lambert and Bilderbeek, 2022). All analyses were run using R version 4.1.0 (R Core Team, 2022). We simulated data under a variety of parameter sets, for an island that is 5 million years old. Mainland extinction rates (*μ*_*M*_) varied between 0 and 2.0, at intervals of 0.2. This is to test a variety of mainland extinction scenarios, although in reality mainland extinction rates are expected to be lower than island extinction rates (Manne et al., 1999; Purvis et al., 2000; Pimm et al., 2014), and theory predicts mainland populations are fitter and thus more resilient to extinction (Rosindell et al., 2015). The mainland pool size was set to *M* = 1000. The mainland sampling fraction was set at 0.5 - 1.0 at 0.1 intervals. When the rate of mainland extinction equals zero and the mainland sampling probability equals one the model reduces to the original DAISIE model. Lower sampling probabilities were not used as it is known that poor sampling negatively impacts the reliability of macroevolutionary inference and thus it is well established that inference should not be conducted on poorly sampled phylogenetic data (Pybus and Harvey, 2000; Cusimano and Renner, 2010). We chose empirically realistic rates for island cladogenesis, extinction, anagenesis and colonisation (*λ*^*c*^ = 0.5, *μ* = 0.25, *λ*^*a*^ = 0.5 and *γ* = 0.01, all rates have units of per species per million years) (Valente et al., 2015, 2017b, 2019). We ran simulations with varying levels of diversity-dependence (*K*^*′*^ = 5, 50). The higher carrying capacity (*K*^*′*^= 50) is a high diversity limit on each island clade, and thus clades will experience little diversity-dependence on the island. In total, we tested 46 parameter combinations, each simulated for 500 replicates to account for stochasticity, with each replicate producing an *ideal*, an *empirical*_*ME*_, an *empirical*_*US*_ and an *empirical*_*UD*_ data set.

We compared the *ideal* simulated data to the *empirical*_*ME*_, *empirical*_*US*_ and *empirical*_*UD*_ simulated data to determine what signatures mainland dynamics and sampling leaves in the data. We examined three metrics in each data set: (1) the difference in normalised cumulative number of island colonisation events through time (comparison using the normalised colonisations through time (ΔnCTT) statistic, see below), (2) the percentage of *maximum island age colonisations* (i.e. cases in which the most recent colonisation time retrieved from the phylogenetic tree of mainland and island species falls before the island age), and (3) the percentage of species on the island that are endemic at the present.

The difference in two nCTT curves (ΔnCTT) is calculated as:

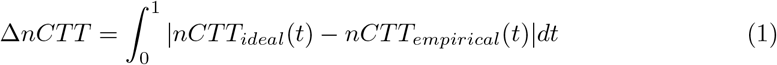

where nCTT is the cumulative number of colonisation events (i.e. of species that survive to the present) through time on the island, normalised for both time and number of species so each are within [0, 1]. *nCTT*_*ideal*_ and *nCTT*_*empirical*_ refer to *ideal* and *empirical* data respectively for the same replicate. This formula is equivalent to the nLTT statistic for quantifying the difference in cumulative number of species between lineage through time plots and uses the nLTT R package version 1.4.5 (Janzen et al., 2015; Janzen and Bilderbeek, 2021).

For each data set, we estimated parameters using the DAISIE maximum likelihood (ML) inference model (see supplementary methods for details), from the DAISIE R package version 4.1.1 (Etienne et al., 2022), and compared the estimates between the *ideal* and *empirical* (*empirical*_*ME*_, *empirical*_*US*_, and *empirical*_*UD*_) data sets. We fitted the diversity-dependent DAISIE model, because all data sets were simulated under diversity-dependence. For each likelihood optimisation the initial parameter estimates were the generating parameter values to help find the global likelihood optimum, because we are interested in errors arising from differences in the data and not arising from imperfect optimisation performance. To determine the error made when applying the DAISIE inference model to the *empirical* data relative to the *ideal* data we compared the estimates from each to the generating values. We expected that even in the *ideal* case the DAISIE inference model would bias rate estimates given that it can only account for endemics produced under evolutionary processes on the island and does not consider any mainland dynamics (e.g., a species becoming endemic not due to speciation on the island but due to extinction on the mainland).

## Results

### Summary of simulated data

Across all parameter sets, the mean total species diversity for the *ideal* and *empirical* data was 74.15 species (max = 199 and min = 19) (Fig. S3, all *empirical* data sets were equal in this metric). The mean number of colonisation events for the *ideal* data was 34.04 (max = 57 and min = 14) (Fig. S3). For the *empirical* data, the mean number of colonisations was 33.69 (max = 55 and min = 14) (Fig. S3). The number of colonisation events differed between *ideal* and *empirical* data sets in a few cases due to some scenarios causing multiple colonisations to be collapsed into a single clade when the full information is not known in the *empirical* case (all *empirical* data sets were equal). Every simulated island data set contained sufficient data (i.e. *>* 5 colonisations and *>* 10 species) to reliably test inference performance of DAISIE, while still being small enough to represent realistic island biodiversity.

### Signatures of mainland extinction on data

The difference in colonisation through time (ΔnCTT) between *ideal* and *empirical*_*ME*_ data increases with mainland extinction but plateaus around 0.15 above a mainland extinction rate of 1.0 (Fig. 3A). Evidently, there are no species that are classified as *maximum island age colonisations* in the *ideal* data, as it is always known at which time a species colonises the island (Fig. 4A). However, in the *empirical*_*ME*_ data even a low rate of mainland extinction (e.g. *μ*_*M*_ = 0.2) causes a jump in the mean percentage of *maximum island age colonisations* to 20% (Fig. 4B), with the median percentage *maximum island age colonisations* plateauing at around 25 - 30%. By contrast, the difference in the percentage of endemic species on the island between *ideal* and *empirical*_*ME*_ data sets is minimal, with both increasing with higher rates of mainland extinction (Fig. 4C,D).

**Figure 3:**
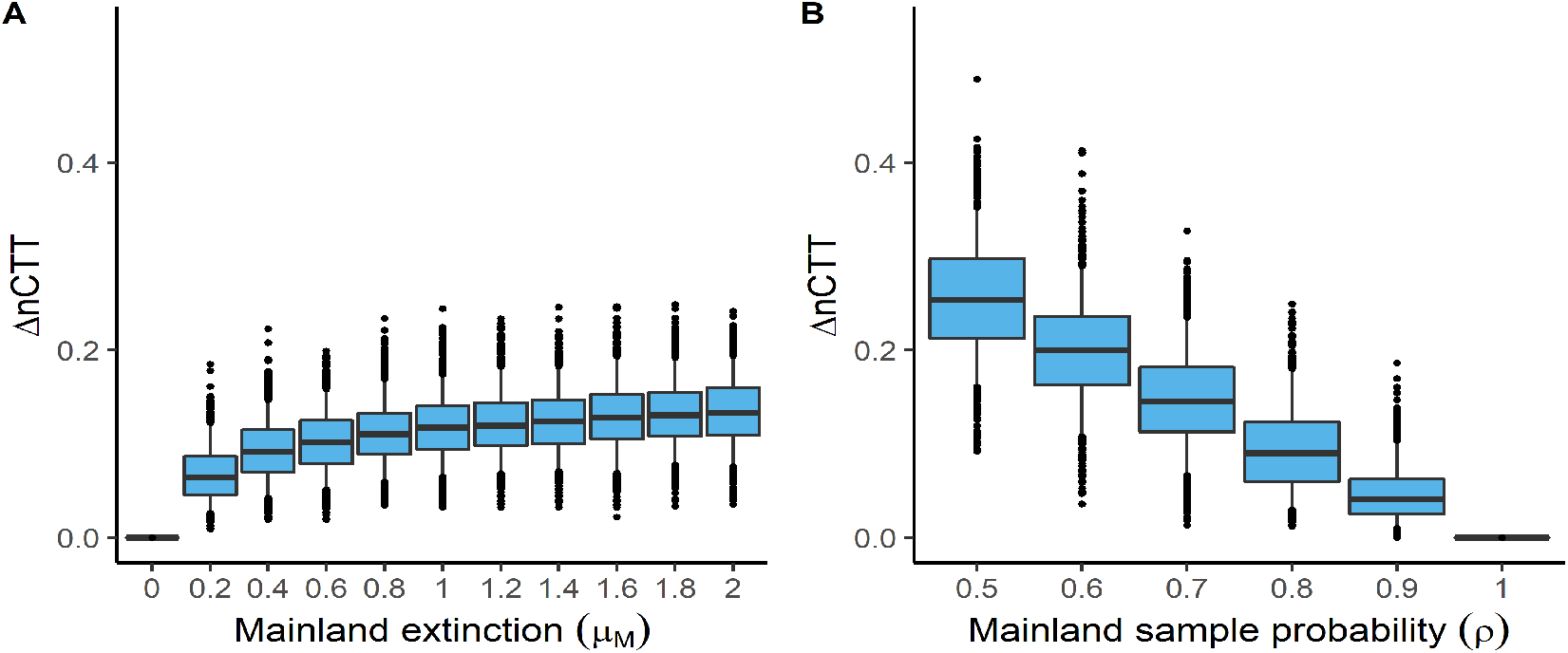
Difference in normalised colonisation through time (ΔnCTT) values for different values of (A) mainland extinction (*μ*_*M*_) and (B) mainland sampling probability (*ρ*). Only the unsampled scenario is shown in this figure because unsampled and undiscovered have identical ΔnCTT values. Boxes show the median, 25th and 75th percentiles, while the whiskers extend to the 5th and 95th percentiles. The dots represent points outside of this range.

**Figure 4:**
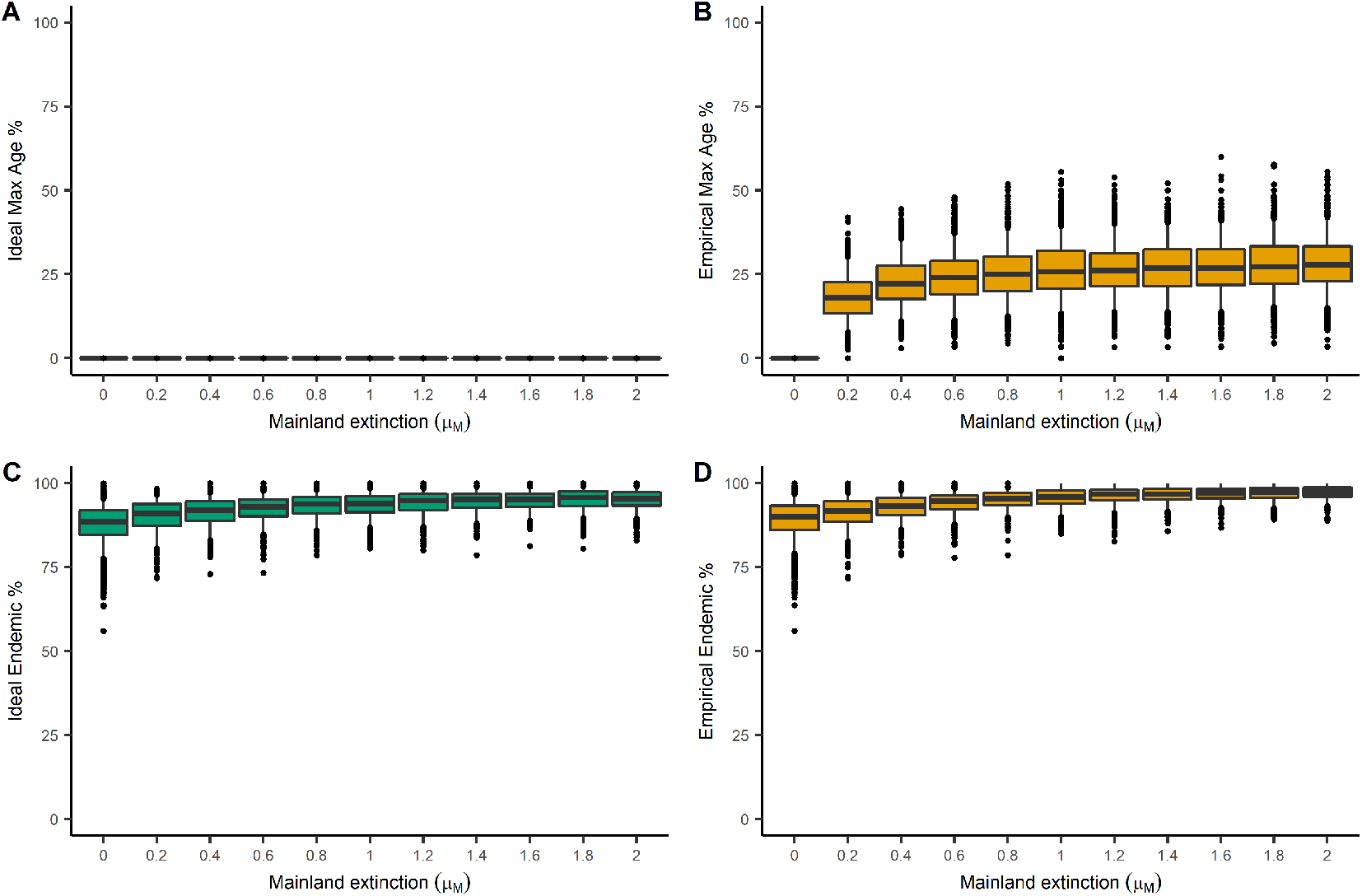
The effect of mainland extinction on *maximum island age colonisations* and endemism. The top two panels show the percentage of *maximum island age colonisations* (Max age) for different values of mainland extinction (*μ*_*M*_), for (A) *ideal* and (B) *empirical*_*ME*_ data. The bottom panels show the percentage of island species that are endemic, for (C) *ideal* and (D) *empirical*_*ME*_ data. Boxes show the median, 25th and 75th percentiles, while the whiskers extend to the 5th and 95th percentiles. The dots represent points outside of this range.

### Signatures of unknown and undiscovered species sampling on data

A decrease in mainland sample probability produces an approximately linear increase in the ΔnCTT (Fig. 3B), in contrast to the asymptotic ΔnCTT produced by increasing mainland extinction rate (Fig. 3A). *Ideal* data contains no *maximum island age colonisations* as is the case for mainland extinction (Fig. S4A, S5A). The percentage of *maximum island age colonisations* increases when the sampling probability of mainland species decreases for both *empirical*_*US*_ and *empirical*_*UD*_ scenarios (Fig S4B, S5B). At *ρ* = 1 there are no *maximum island age colonisations* as all mainland species are sampled. Island endemism is approxi-mately equal across sampling probabilities in the *ideal, empirical*_*US*_ and *empirical*_*UD*_ data (Fig S4C,D, S5C,D). Sampling does not affect the *ideal* data, ergo, there is no major differences in the pattern of percentage of island endemic species for unsampled and undiscovered sampling (Fig S4C, S5C).

### Inference error from mainland dynamics

When there is no mainland extinction and mainland sampling is complete, the *ideal* and *empirical*_*ME*_ data are identical. As a result, the parameter estimates of the two data sets are also identical (Fig. S6). The parameter values estimated are clustered around the true value used to simulate the data, with only few estimates being far from the true value, confirming DAISIE’s good performance when assumptions of the inference model are met. For non-zero mainland extinction rates, the difference in estimation between the *ideal* and *empirical*_*ME*_ is already visible from the lowest rate of mainland extinction (*μ*_*M*_ = 0.2, Fig. 5), and increases with larger mainland extinction rate (Fig. 6). Still, estimated rates of cladogenesis, extinction and colonisation are spread around the true values for *ideal* data, whereas anagenesis is relatively more biased (Fig 5, 6). This is consistent with the expectation of an over-estimation in the inference of anagenesis from *ideal* and *empirical*_*ME*_ data, with *empirical*_*ME*_ being more biased (Fig. 5, 6).

**Figure 5:**
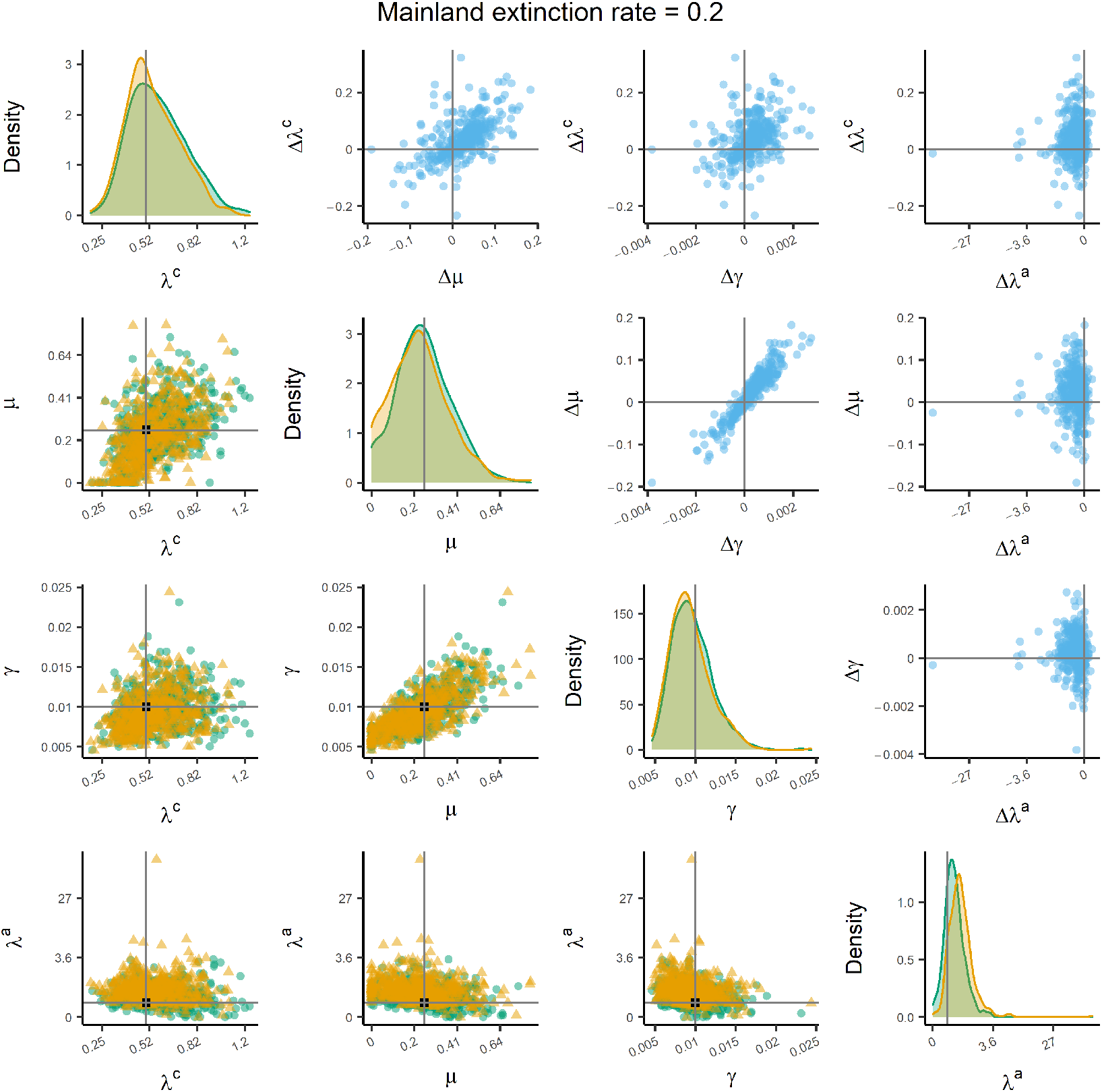
Parameter estimates inferred from the *ideal* and *empirical*_*ME*_ data sets of a simulation with a mainland extinction rate of 0.2. The diagonal panels show the distributions of the parameter estimates from the *ideal* (green) and *empirical*_*ME*_ (yellow) data sets. The vertical grey line is the true value used to simulate the data. The panels on the lower triangular show the point estimates from the *ideal* (green circles) and *empirical*_*ME*_ (yellow triangles) data sets. The black square at the intersection of the grey lines is the true value used to simulate the data. The panels on the upper triangular show the differences between the green circles and yellow triangles (i.e. *ideal* minus *empirical*_*ME*_). The grey lines intersect at zero. The *x* -axis on the diagonal panels and both axes on the off-diagonal panels are transformed with inverse hyperbolic sine transformation (ln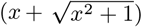) to display data (including zeros and negative values) over several orders of magnitude; the tick labels present the untransformed values. These results were simulated with *K*^*′*^ = 5 and are qualitatively similar to the results from *K*^*′*^ = 50. Anagenesis (*λ*^*a*^) is fixed under certain conditions (see supplementary methods).

**Figure 6:**
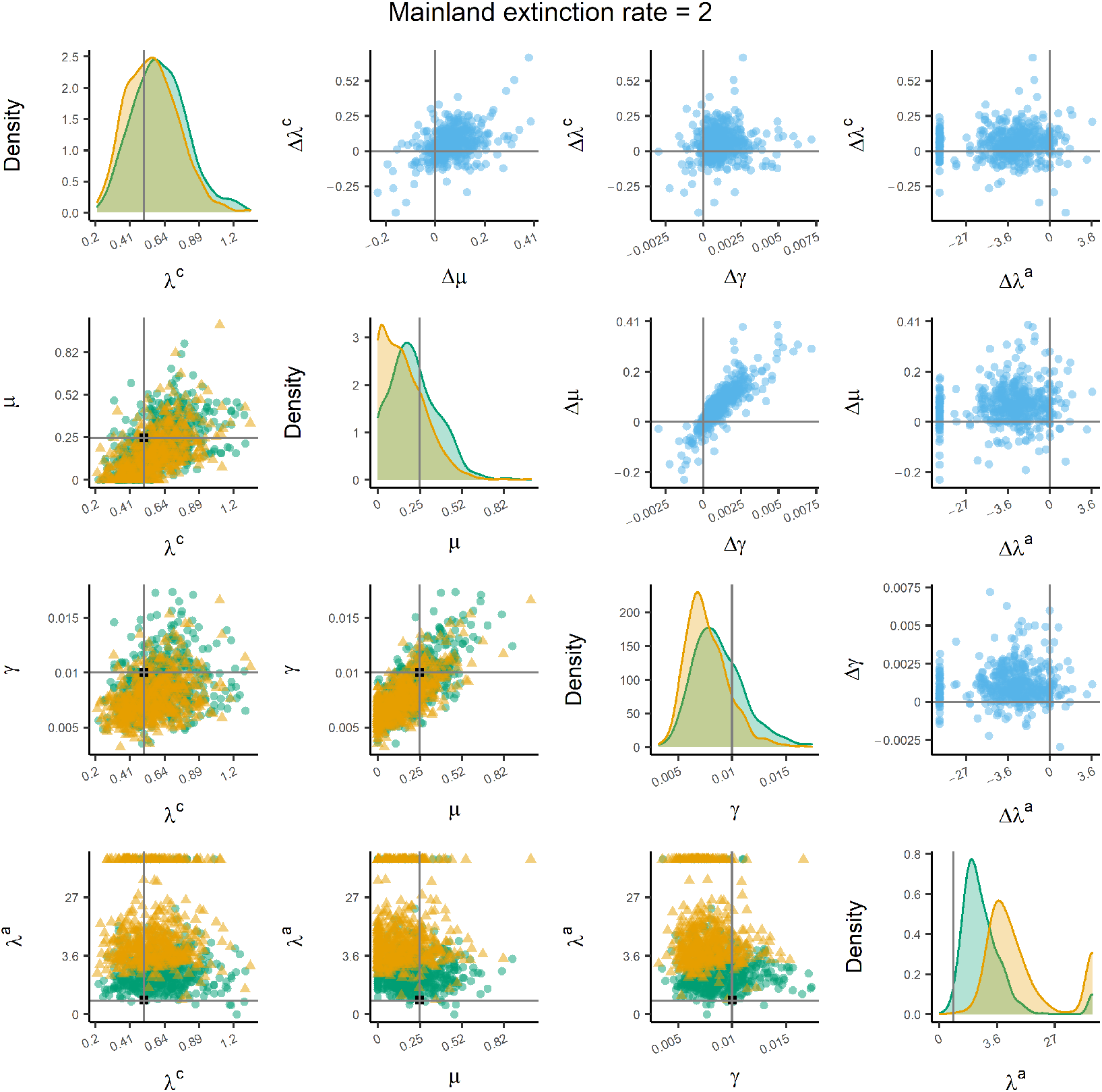
Parameter estimates inferred from the *ideal* and *empirical*_*ME*_ data sets of a simulation with a mainland extinction rate of 2.0. The diagonal panels show the distributions of the parameter estimates from the *ideal* (green) and *empirical*_*ME*_ (yellow) data sets. The vertical grey line is the true value used to simulate the data. The panels on the lower triangular show the point estimates from the *ideal* (green circles) and *empirical*_*ME*_ (yellow triangles) data sets. The black square at the intersection of the grey lines is the true value used to simulate the data. The panels on the upper triangular show the differences between the green circles and yellow triangles (i.e. *ideal* minus *empirical*_*ME*_). The grey lines intersect at zero. The *x* -axis on the diagonal panels and both axes on the off-diagonal panels are transformed with inverse hyperbolic sine transformation (ln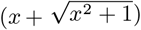) to display data (including zeros and negative values) over several orders of magnitude; the tick labels present the untransformed values. These results were simulated with *K*^*′*^ = 5 and are qualitatively similar to the results from *K*^*′*^ = 50. Anagenesis (*λ*^*a*^) is fixed under certain conditions (see supplementary methods).

Parameter estimation for colonisation and diversification (cladogenesis and extinction) was reliable for the *ideal* data but slightly negatively biased for *empirical* data with high levels of undiscovered species (Fig. S7). In the case of undiscovered species, the anagenesis rate was positively biased, because of the higher island endemicity, with the bias being pronounced when a large proportion of the mainland species are undiscovered (Fig. S7, S8). The bias in all rates was negligible in the unsampled mainland species case, because although the colonisation time can be biased by the sampling, the endemicity status is correctly assigned and thus the DAISIE inference model does not overcompensate by elevating anagenesis (Fig. S9, S10).

Mainland speciation and extinction, and as a result the non-independence and interference of species on the island from the same mainland clade, does not influence the estimates of a clade-specific carrying capacity (Fig. 7). This is likely because island colonisation is a rare event and thus the chance that two colonisations derive from the same mainland clade is small. The estimated carrying capacity is approximately equal across all levels of mainland extinction rate (Fig. 7). The percent of carrying capacities that were estimated to be infinite did not show a dramatic change across rates of mainland extinction, but did show a slight negative trend (Fig. S11). The estimates of carrying capacity are similar for *ideal* and *empirical*_*ME*_ estimates. Across the parameter space the mean estimate of the carrying capacity is below the true value, the upper quantile is around the true value, and there are a few estimates that are very large (Fig. 7). The carrying capacity is well estimated in the unsampled and undiscovered species scenarios, but this is expected as mainland species are independent and do not undergo speciation or extinction in these scenarios (Fig. S12, S13).

**Figure 7:**
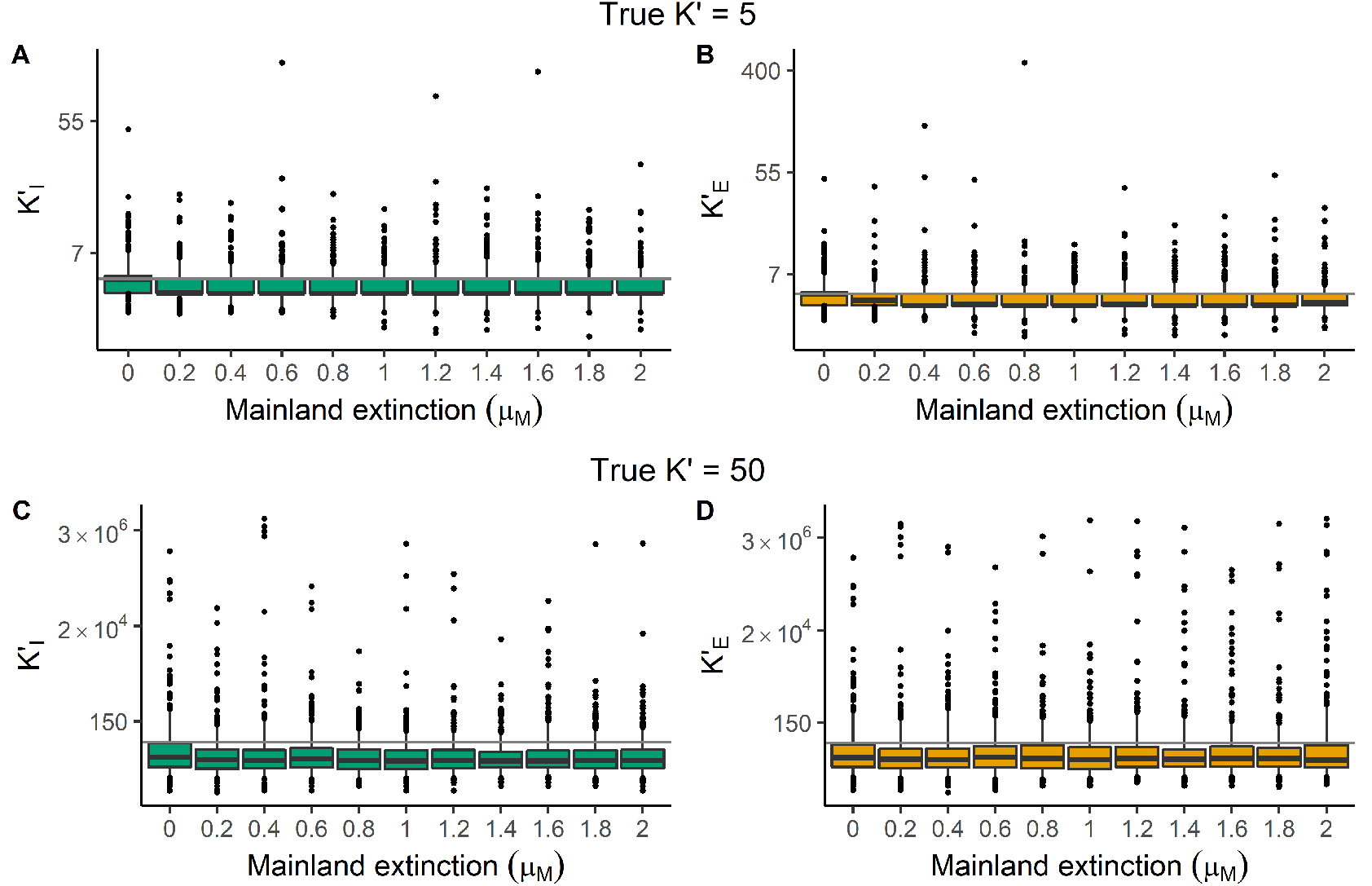
Estimates of carrying capacity parameter (*K*^*′*^) for (A, C) *ideal* data sets (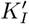, green) and (B, D) *empirical*_*ME*_ data sets (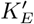, yellow) under two simulated carrying capacities *K*^*′*^ = 5 (A, B) and *K*^*′*^ = 50 (C, D), plotted for different values of mainland extinction (*μ*_*M*_). The grey horizontal line is the true value. Boxes show the median, 25th and 75th percentiles, while the whiskers extend to the 5th and 95th percentiles. The dots represent points outside of this range. The *y* -axis is plotted on a natural log scale to display data over several orders of magnitude.

## Discussion

Macroevolutionary studies of islands typically focus on insular taxa, sampling only a few species on the mainland that are thought to be close relatives or ancestors of the target islands species, and largely ignoring diversification dynamics on the continent. But how much can we infer about island evolution without a good understanding and sampling of the mainland? The DAISIE inference model uses phylogenetic data from an island community to estimate rates of colonisation, speciation and extinction, and has been shown to do so quite accurately (Valente et al., 2017a, 2018), but this accuracy is predicated on the assumption that the mainland species pool is composed of independent static species, and all mainland species are sampled and known. Here we show that mainland dynamics – speciation, extinction and phylogenetic relatedness – and incomplete sampling leave clear signatures in *empirical* data, with a higher proportion of endemic species and colonisation times further in the past, sometimes exceeding the age of the island, when compared to data for which there is complete information on all the mainland species. Nevertheless, even in the presence of mainland extinction and incomplete mainland sampling, DAISIE - which ignores these mainland processes - can generally still estimate rates and dynamics of island colonisation and diversification reliably. Mainland species that have gone extinct before the present or are undiscovered cause an overestimation of anagenesis, which we attribute to the higher perceived island endemicity. Therefore, although the study of macroevolutionary dynamics is constrained by the trade-off between model complexity and data availability, with complete – fossil and molecular – data improving phylogenetic reconstruction and hence the macroevolutionary rates estimates (Silvestro et al., 2014; Zhang et al., 2016), we show that even when applying a model to incomplete data the inferred parameter estimates are close to the true values, and thus the model is adequate and robust. This robustness to violations in the inference model’s assumptions is congruent with previously confirmed robustness to violation in the assumption of constant island area (Neves et al., 2021). Even though there may be fundamental limits to phylogenetic inference (Losos, 2011), when DAISIE is applied correctly to test hypotheses in island biogeography, it can be a useful tool (Morlon et al., 2020).

The influence of mainland dynamics did not vastly differ between low and high rates of mainland extinction. The reliable performance of DAISIE for different mainland dynamics was shown by the variance in the estimates of cladogenesis, extinction and colonisation rates remaining roughly equal across different levels of mainland extinction. The generally accurate estimation of these rates is in contrast to the expectation that they would be un-derestimated due to the colonisation time being biased back in time. A logical follow-up question is what the mainland rates are in nature, both in absolute terms and relative to rates on the island. Islands are theorised as potential species sinks (i.e. high extinction rate) or as ideal settings for species radiations (i.e. high speciation rate). It is likely that continental regions have lower extinction rates than islands and thus the high rate of mainland extinction used in this study (up to four times higher than island extinction) are empirically improbable (Manne et al., 1999; Purvis et al., 2000; Pimm et al., 2014), but even with these unrealistically high rates, the inference model is quite robust. The few studies looking into island-mainland dynamics have found that island diversification rate (speciation rate minus extinction rate) is similar between the two systems (Algar and Losos, 2011; Román-Palacios and Wiens, 2018; Patton et al., 2021), with extinction rates close to zero, likely due to common issues in estimating extinction from reconstructed trees (Etienne et al., 2012; Louca and Pennell, 2021). Under the assumption that biodiversity is in equilibrium on the mainland the extinction rate could be approximated by the speciation rate, which can be relatively well-inferred (Marshall, 2017).

At a relatively low rate of mainland extinction some estimates of anagenesis were large, and as the rate of mainland extinction increases, the number of large estimates becomes more numerous. Anagenesis is commonly interpreted as the process by which island and mainland populations diverge and speciate in isolation, either due to selective or stochastic processes. Although the estimates of the rate of anagenesis were higher than the true value, especially in the *empirical* data, this parameter can be reinterpreted as the signal of island species becoming endemic either through divergence of extant populations or through species turnover on the mainland. Mainland extinction and incomplete mainland data produced similar patterns in the island data and inference, but showed a few differences. The most obvious is the different level of bias in anagenesis, with it being overestimated for undiscovered species and reliably estimated for incomplete sampling due to unsampled known species. This is in line with expectations, because in the case of undiscovered mainland species we erroneously assign the endemic status to an island species, whereas in the case of an unsampled mainland species we do not.

In light of the signatures mainland dynamics and incomplete sampling leave in the data, it seems previous analyses using DAISIE were not severely affected, as the majority of colonisation events fell within the existence of the island, and many archipelagos had several non-endemic species (i.e Valente et al. (2020)). The level of bias in colonisation times in previous applications of the model is unknown. However, given our findings the results of previous studies seem to be qualitatively robust, but may have minor errors in estimated rates, with non-random extinction being a potential avenue for future analysis. For example, one interesting case study is that of birds from the archipelagos of Macaronesia. On the Azores, the Canary Islands, and Madeira most colonisation times have been estimated using phylogenetic data to be close to the present, while a few are *maximum island age colonisations* (Valente et al., 2017b, 2020). This may indicate non-random extinction on the mainland with some clades experiencing greater error in colonisation times than others. By contrast, all colonisations of Cape Verde were estimated to have occurred near the present. Cape Verde has a mainland source pool further south than the other Macaronesian archipelagos (off the coast of Senegal), suggesting possible geographical variation in extinction rates on the mainland. Thus, it is likely that taxonomic and global variation in mainland extinction rates will lead to different levels of impact for different island systems. Furthermore, the effect of mainland extinction will depend on the topology and branching structure of the phylogeny, analogous to how evolutionary history loss depends on topology, if extinction happens on long branches the effect will be worsened (Nee and May, 1997).

In this study we show that there is minimal error produced in the estimation of island rates due to mainland dynamics when the empiricist has access to multiple samples from each species (i.e. at least one sample from the mainland and the island for non-endemic species), providing sufficient resolution in the phylogenetic tree to identify colonisation times. However, in cases where phylogenetic data only has a single sample per species, as often occurs in practice, the colonisation times will be further from the true value. For example, empiricists extracting colonisation times from phylogenies at the species-level (e.g. bird (Jetz et al., 2012) or mammal (Upham et al., 2019) phylogenies) should be more cautious about the reliability of their estimates. Furthermore, mainland extinction and incomplete mainland sampling are only two mechanisms that potentially bias island biogeographical data, but there are other sources of bias that we did not investigate but may influence the data and inference. For instance, the divergence time is often inferred from the coalescence time of genetic sequences, but this can predate the speciation or colonisation time (Kingman, 1982). This can bias phylogenetic data for island clades by pushing the speciation or colonisation time further in the past, most likely from ‘deep coalescence’ from incomplete lineage sorting (Maddison, 1997), especially for large mainland effective population sizes or recent island colonisations. Lastly, the data simulated here provides exact branching times (i.e. without error), whereas empirical phylogenetic trees have uncertainty in branching times (Kapli et al., 2020).

Some of our results could be artifacts from the design of our simulation of mainland dynamics. Due to the use of a Moran process, the increase in mainland extinction is intrinsically paired with the rate of mainland speciation. Thus, the plateau in error (Fig. 3A, 4) may occur because the error in colonisation time estimation due to extinction is reduced by frequent branching events stopping the colonisation time from jumping back to the island age. Conversely, in the case of zero mainland extinction, under the Moran process mainland clades do not form because each mainland lineage starts as a single species and can only diversify once another species on the mainland goes extinct. When incomplete sampling is applied to these monotypic lineages, the colonisation is pushed back to the island age, which is probably not representative of continental phylogenies, unless the island is very young. The use of a birth-death process instead of the Moran process would, however, have its own limitations, namely, when speciation exceeds extinction (*λ > μ*) the mainland species pool is expected to grow exponentially through time, and hence one would not be able to tell whether any differences result from extinction dynamics or from non-constant mainland pool size. Moreover, it is not in line with evidence that mainland diversification is not a constant-rate birth-death process, but more likely a diversity-or time-dependent process (Condamine et al., 2019; Henao Diaz et al., 2019).

This study used a novel simulation model instead of developing a novel likelihood incorporating island and mainland dynamics. Developing a likelihood is feasible if the dynamics on the mainland are precisely known, i.e. if a phylogenetic tree is available for all species, including now-extinct ones, that can have immigrated to the island. For each species on this tree, it is precisely known when it was present on the mainland and for this period the current DAISIE likelihood for each colonising clade can be used. However, in practice the phylogenetic tree will contain only extant species, and a likelihood requires integrating over all possible complete trees consistent with this extant-species-only tree. There are methods to do this, for example using data augmentation to simulate the missing data and then applying expectation maximization (EM) (Richter et al., 2020), but this is far from trivial. However, even a complete phylogeny of the mainland with only extant species is often not available in practice, particularly if the taxonomic scope of the analysis is wide. With the new simulation described in this study, a simulation-based inference, such as Approximate Bayesian Computation (ABC), could be used, but ABC is probably not feasible for estimation in this case because the dimensionality of the model would require a computationally prohibitive number of simulations. Our study shows that all these efforts are not needed in most cases. Thus, robustness studies such as this are useful to explore what aspects are important for future development of inference methods.

It remains crucial that model development is paired with an understanding of model performance. Models never encapsulate the complexity of species diversification, but once the ‘danger zones’ of model performance are elucidated (e.g. Felsenstein zone (Huelsenbeck and Hillis, 1993), and anomalous gene trees (Degnan and Rosenberg, 2006), in phylogenetic inference), and empirical indistinguishability (for birth-death models, (Louca and Pennell, 2020), and species-abundance distributions for neutral models, (Haegeman and Etienne, 2011)), appropriate application can be carried out. Due to the deep timescales at which it often operates, island biogeography is a field that requires inference models to understand its largest open questions (Patinõ et al., 2017), so as novel data becomes available, knowing which models can be applied when is crucial. We hope to have taken an important step in doing so for island biogeography models.

## Acknowledgments

We would like to thank the Theoretical and Evolutionary Community Ecology group at the University of Groningen, Pratik Rajan Gupte and Shu Xie for discussions and their input. We would like to thank the Center for Information Technology of the University of Groningen for their support and for providing access to the Peregrine high performance computing cluster. JWL was funded through a Study Abroad Studentship by the Leverhulme Trust and was also funded by a NWO VICI grant awarded to RSE who was funded by the same grant. PSN was funded through a FCT PhD Studentship with reference SFRH/BD/129533/2017, co-funded by the Portuguese Ministério da Ciência, Tecnologia e Ensino Superior and the European Social Fund. LV was funded by a NWO VIDI grant.

## Supplementary Material

### Supplementary Methods Total rates

Here we provide the formulas used for the processes on the island.

Cladogenesis:

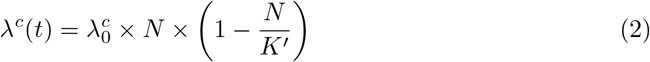

Extinction:

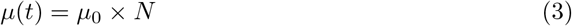

Colonisation:

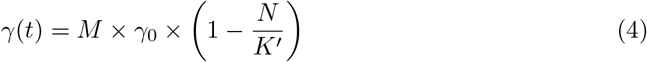

Anagenesis:

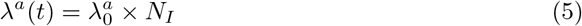

Mainland extinction:

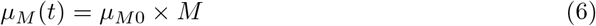

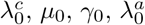 are the intrinsic rates of cladogenesis, extinction, colonisation, and anagenesis respectively. *N* is the number of species on the island at time *t. K*^*′*^ is the carrying capacity of an island clade. *M* is the number of mainland species at time *t. N*_*I*_ is the number of immigrant species (non-endemic species) on the island at time *t. μ*_*M*0_ is the intrinsic mainland extinction rate.

### DAISIE inference

When fitting the DAISIE maximum likelihood model to data we optimised all parameters of the model (*λ*^*c*^, *μ, K*^*′*^, *γ, λ*^*a*^). There was one exception to this, in the case that all species on the simulated island were endemic and there are no recolonisations. In this case we already know beforehand the maximum likelihood estimate for the anagenesis rate, as we explain below. There are two scenarios when all island species are endemic. First, if all island colonists form a clade with more than one species there is no information in the data on anagenesis and it is zero because cladogenesis can explain all the endemic species. Alternatively, if there are some singleton colonists that have not diversified on the island, the anagenesis rate will be infinite, because an infinite rate of anagenesis makes observing all species on the island being endemic most likely. When optimising the likelihood function for a data set in which all island species are endemic (without any recolonists), anagenesis becomes large relative to the other processes, causing the likelihood master equation to become stiff. In solving this stiff equation the integration — using the Runge-Kutta-Fehlberg method — takes many small steps. Taking many small steps can require very large amounts of computational time to solve these equations. Therefore, in the case that all island species are endemic (with some singletons) the anagenesis rate is fixed to 100, with all other parameters being freely optimised. In the case all island species are endemic clades with more than one species, the anagenesis rate is fixed to zero. This prevents the likelihood optimisation failing, either due to time or numerical instability of the solver, and gives anagenesis a more realistic value.

The exception to the two scenarios explained above (all endemics either with or without singleton lineages) is when one or more of the species in the data set are recolonists of the same mainland species that already speciated (either by cladogenesis or anagenesis) on the island. When there are recolonisation events in the data set anagenesis can be optimised. This is because there is information on the anagenesis rate. This information comes from the fact that the colonisation rate is bound by the number of colonisations seen in the data set and cladogenesis is bound by the number of branching events in the data set. Therefore, the colonisation rate and cladogenesis rate cannot be freely optimised to explain the endemic recolonist and thus there is information on the anagenesis rate in order to explain the recolonist.

### Supplementary Results

**Figure S1:**
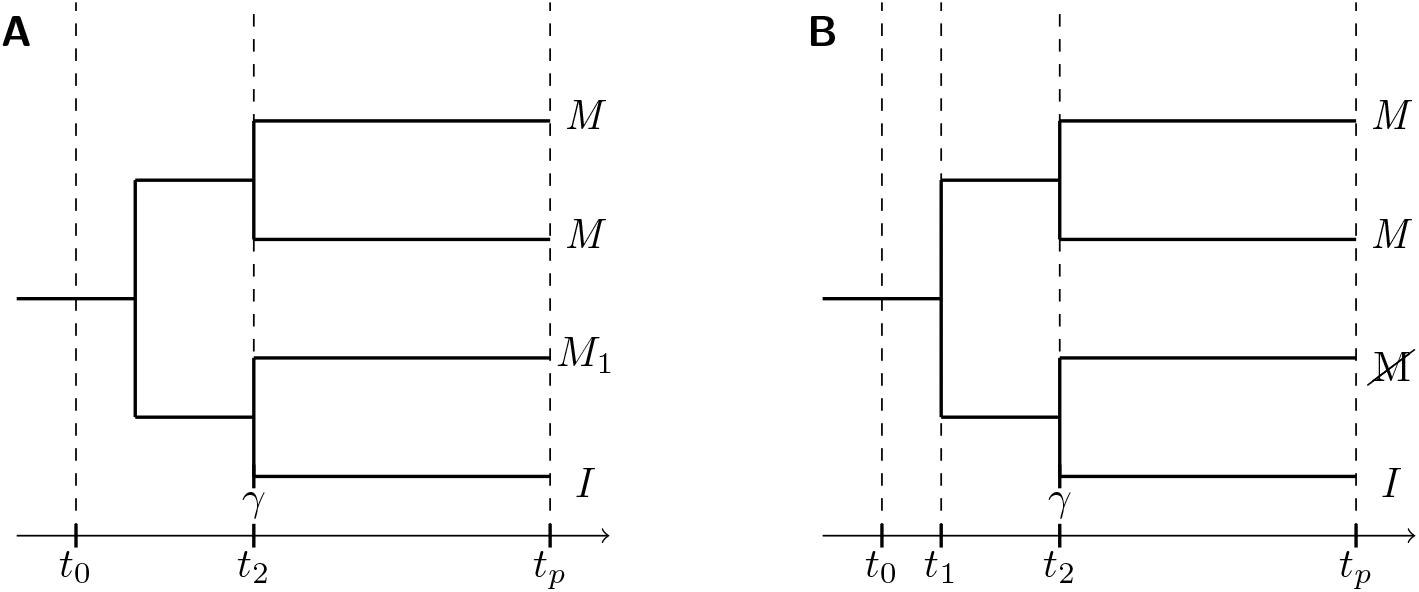
The effect of incomplete sampling is analogous to the effect of mainland extinction for colonisation times (see figure 1 in main text). (A) The colonisation time from the divergence time of the island species (*I*) with its most closely related mainland species (*M*_1_). This approximation overestimates the colonisation time from reconstructed phylogenetic data. The island emerges at *t*_0_, a species colonises the island at *t*_2_ (*γ*) causing divergence from the mainland sister species. *t*_*p*_ is the present. (B) The colonisation time is overestimated when incomplete sampling occurs. Species represented at the tip with 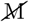 is an extant species on the mainland that is not sampled in the phylogenetic tree. The colonisation of a mainland species to the island occurs at *t*_2_, but as the sister species of the colonist is not sampled in the phylogenetic tree, the colonisation time taken from divergence from a sister species is at *t*_1_.

**Figure S2:**
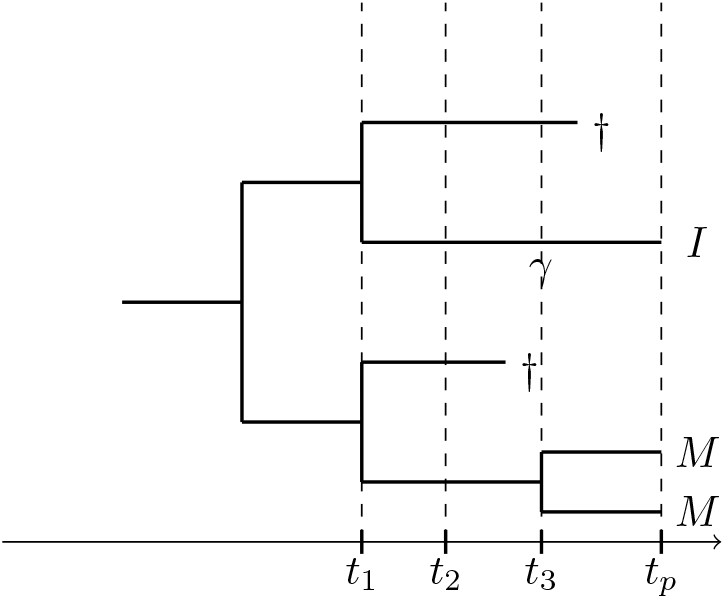
Extinction of species that branched before the emergence of the island does not influence the colonisation times taken from phylogenies. The phylogeny represents a mainland clade, with one species immigrating to the island at *t*_3_, denoted by *γ*. The extant mainland species are denoted by *M* and the island species is denoted by *I. t*_2_ is the time at which the island emerges (it is assumed to be known without error). The timing of colonisation from the phylogeny would be *t*_1_, but the maximum age of the island is *t*_2_, and thus the species cannot have immigrated before *t*_2_. Therefore, the colonisation time is set to have a maximum of *t*_2_. In DAISIE, if the colonisation time extracted from the phylogeny is found to be older than the island, the clade is treated as a *maximum island age colonisation* and the DAISIE inference model integrates across the time from the present to the island’s origin as a possible colonisation time.

**Figure S3:**
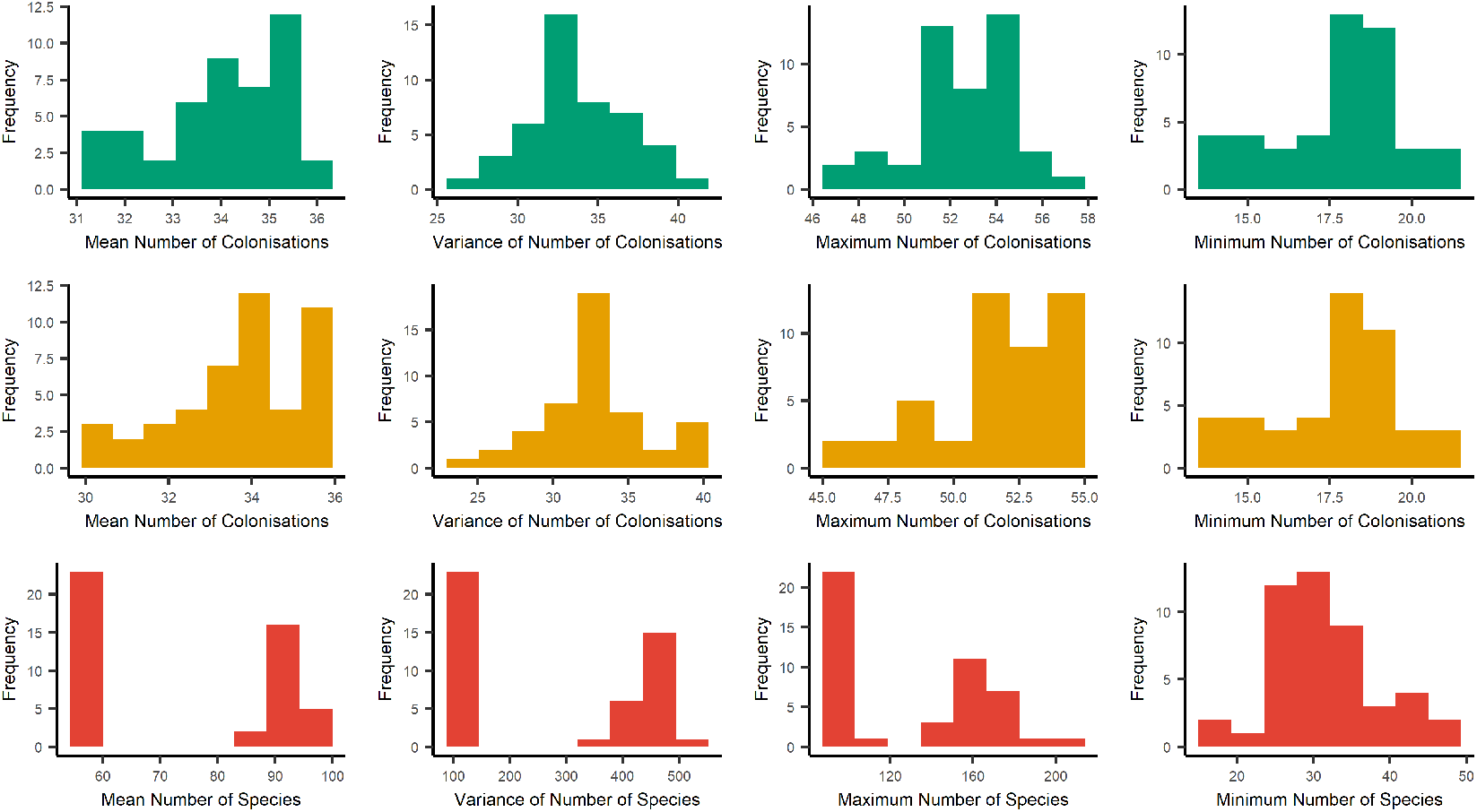
Summary statistics of the simulated data for number of colonisations that survived to the present and number of species on the island at the present. The data is summarised by mean (first column), variance (second column), maximum (third column) and minimum (fourth column). The first row and second row is number of island colonists for the *ideal* data and *empirical* data, respectively (all three *empirical* data sets were equal). The third row is the number of species on the island, all *ideal* and *empirical* data sets were equal for this metric.

**Figure S4:**
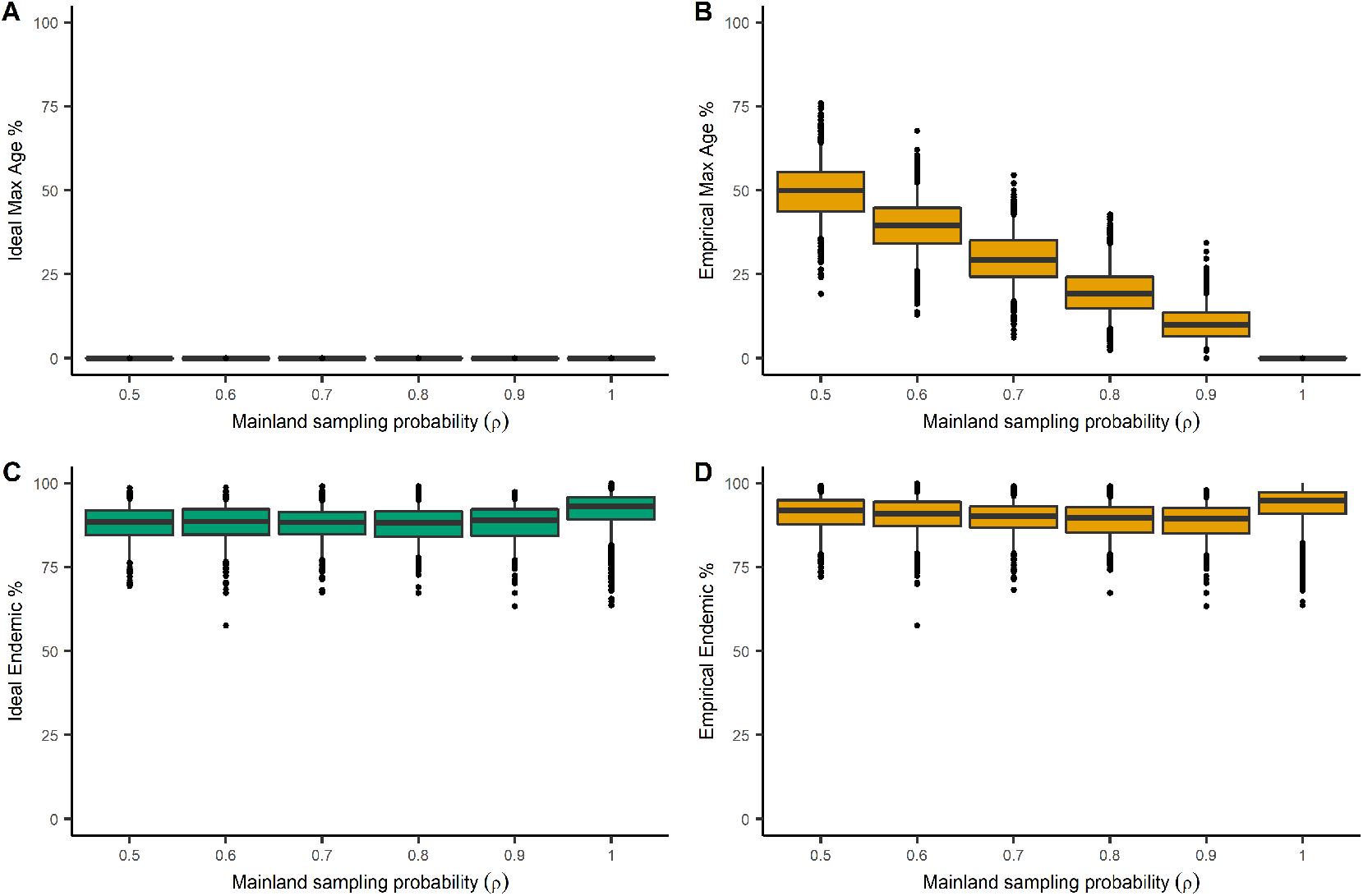
Percentage of *maximum island age colonisation* (max age) for (A) *ideal* and (B) *empirical*_*US*_ data for different values of mainland sampling probability (*ρ*). Percentage of island species that are endemic for (C) *ideal* and (D) *empirical*_*US*_ data. Boxes show the median, 25th and 75th percentiles, while the whiskers extend to the 5th and 95th percentiles, except for anomalous points plotted individually which are defined as greater than the 95th percentile or less than the 5th percentile.

**Figure S5:**
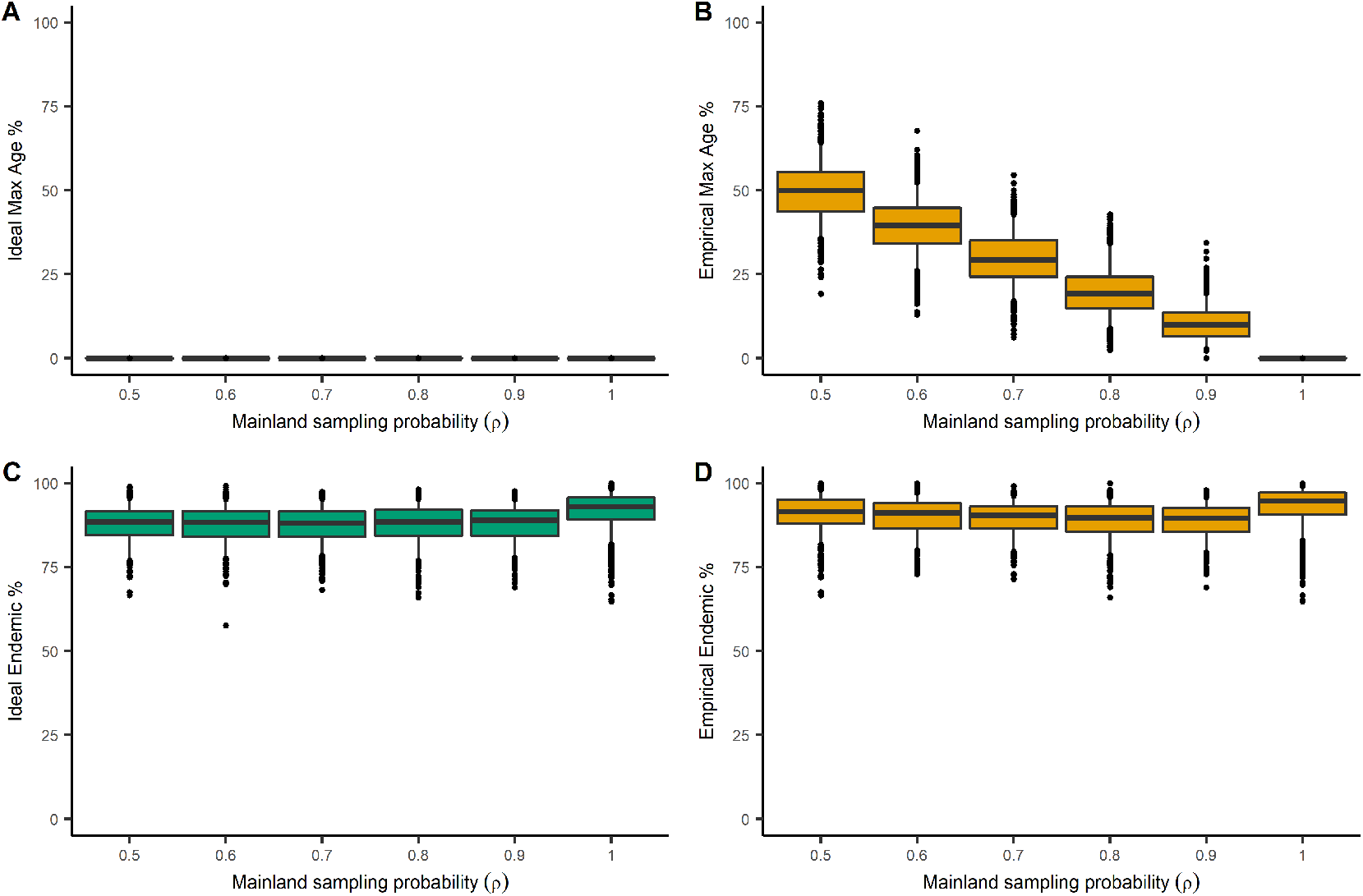
Percentage of *maximum island age colonisation* (max age) for (A) *ideal* and (B) *empirical*_*UD*_ data for different values of mainland sampling probability (*ρ*). Percentage of island species that are endemic for (C) *ideal* and (D) *empirical*_*UD*_ data. Boxes show the median, 25th and 75th percentiles, while the whiskers extend to the 5th and 95th percentiles, except for anomalous points plotted individually which are defined as greater than the 95th percentile or less than the 5th percentile.

**Figure S6:**
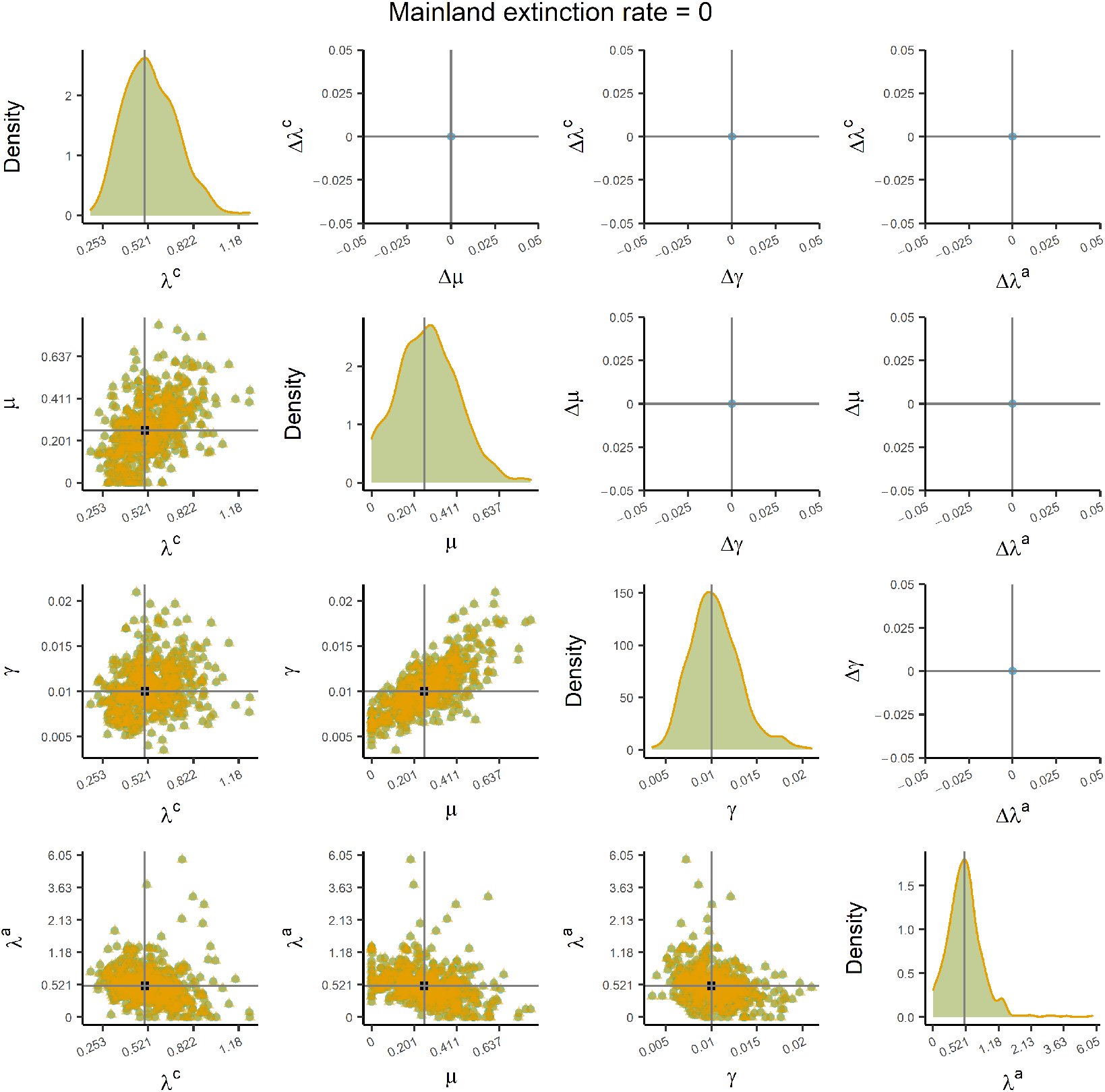
Parameter estimates inferred from the *ideal* and *empirical*_*ME*_ data sets of a simulation with a mainland extinction rate of 0. The diagonal panels show the distributions of the parameter estimates from the *ideal* (green) and *empirical*_*ME*_ (yellow) data sets. The vertical grey line is the true value used to simulate the data. The panels on the lower triangular show the point estimates from the *ideal* (green circles) and *empirical*_*ME*_ (yellow triangles) data sets. The black square at the intersection of the grey lines is the true value used to simulate the data. The panels on the upper triangular show the differences between the green circles and yellow triangles (i.e. *ideal* minus *empirical*_*ME*_). The grey lines intersect at zero. The *x* -axis on the diagonal panels and both axes on the off-diagonal panels are transformed with inverse hyperbolic sine transformation (ln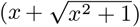) to display data (including zeros and negative values) over several orders of magnitude; the tick labels present the untransformed values. These results were simulated with *K*^*′*^ = 5 and are qualitatively similar to the results from *K*^*′*^ = 50. Anagenesis (*λ*^*a*^) is fixed under certain conditions (see supplementary methods).

**Figure S7:**
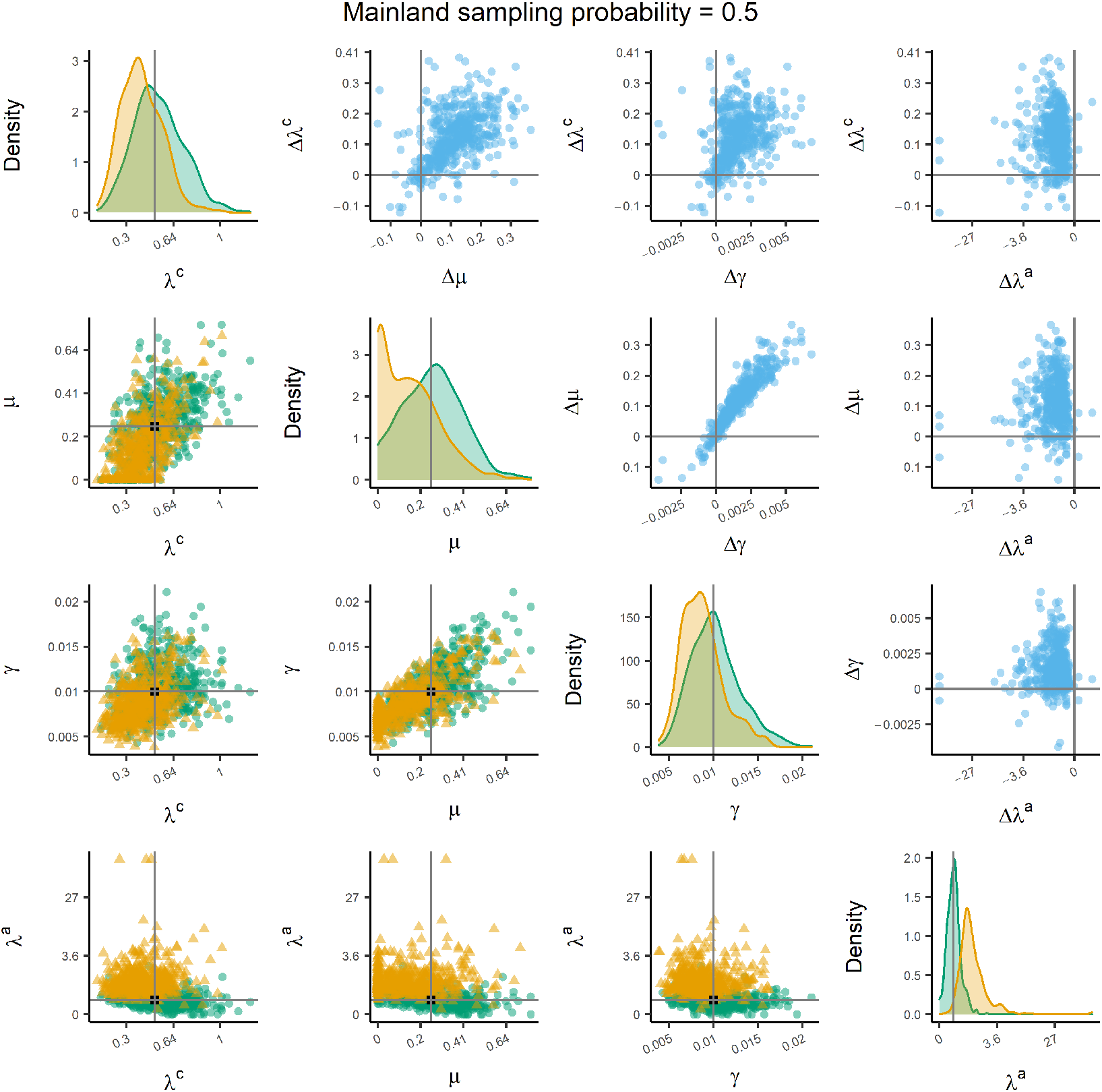
Parameter estimates inferred from the *ideal* and *empirical*_*UD*_ data sets of a simulation with a mainland sampling probability of 0.5. The diagonal panels show the distributions of the parameter estimates from the *ideal* (green) and *empirical*_*UD*_ (yellow) data sets. The vertical grey line is the true value used to simulate the data. The panels on the lower triangular show the point estimates from the *ideal* (green circles) and *empirical*_*UD*_ (yellow triangles) data sets. The black square at the intersection of the grey lines is the true value used to simulate the data. The panels on the upper triangular show the differences between the green circles and yellow triangles (i.e. *ideal* minus *empirical*_*UD*_). The grey lines intersect at zero. The *x* -axis on the diagonal panels and both axes on the off-diagonal panels are transformed with inverse hyperbolic sine transformation (ln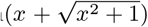) to display data (including zeros and negative values) over several orders of magnitude; the tick labels present the untransformed values. These results were simulated with *K*^*′*^ = 5 and are qualitatively similar to the results from *K*^*′*^ = 50. Anagenesis (*λ*^*a*^) is fixed under certain conditions (see supplementary methods).

**Figure S8:**
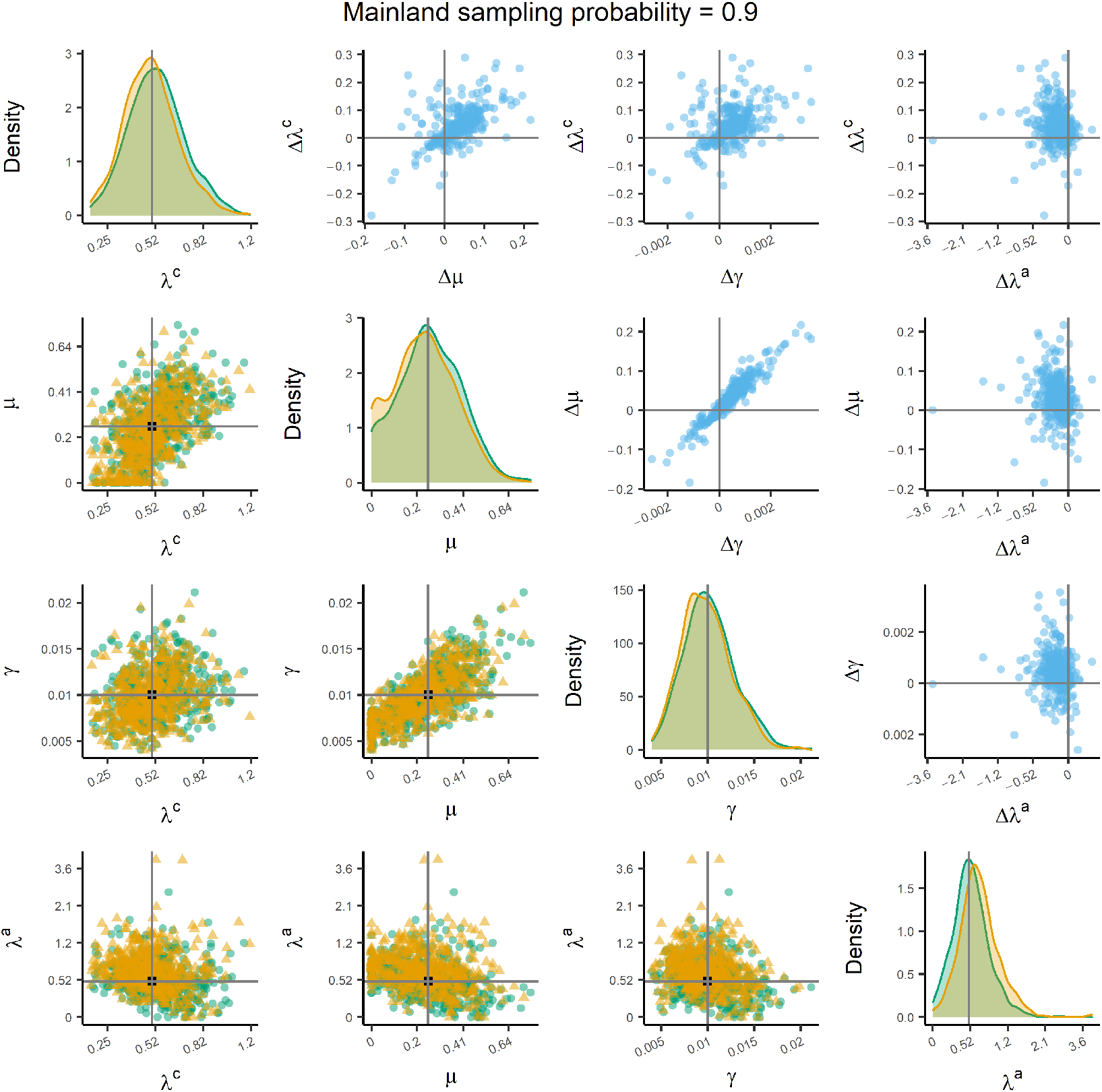
Parameter estimates inferred from the *ideal* and *empirical*_*UD*_ data sets of a simulation with a mainland sampling probability of 0.9. The diagonal panels show the distributions of the parameter estimates from the *ideal* (green) and *empirical*_*UD*_ (yellow) data sets. The vertical grey line is the true value used to simulate the data. The panels on the lower triangular show the point estimates from the *ideal* (green circles) and *empirical*_*UD*_ (yellow triangles) data sets. The black square at the intersection of the grey lines is the true value used to simulate the data. The panels on the upper triangular show the differences between the green circles and yellow triangles (i.e. *ideal* minus *empirical*_*UD*_). The grey lines intersect at zero. The *x* -axis on the diagonal panels and both axes on the off-diagonal panels are transformed with inverse hyperbolic sine transformation (ln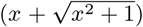) to display data (including zeros and negative values) over several orders of magnitude; the tick labels present the untransformed values. These results were simulated with *K*^*′*^ = 5 and are qualitatively similar to the results from *K*^*′*^ = 50. Anagenesis (*λ*^*a*^) is fixed under certain conditions (see supplementary methods).

**Figure S9:**
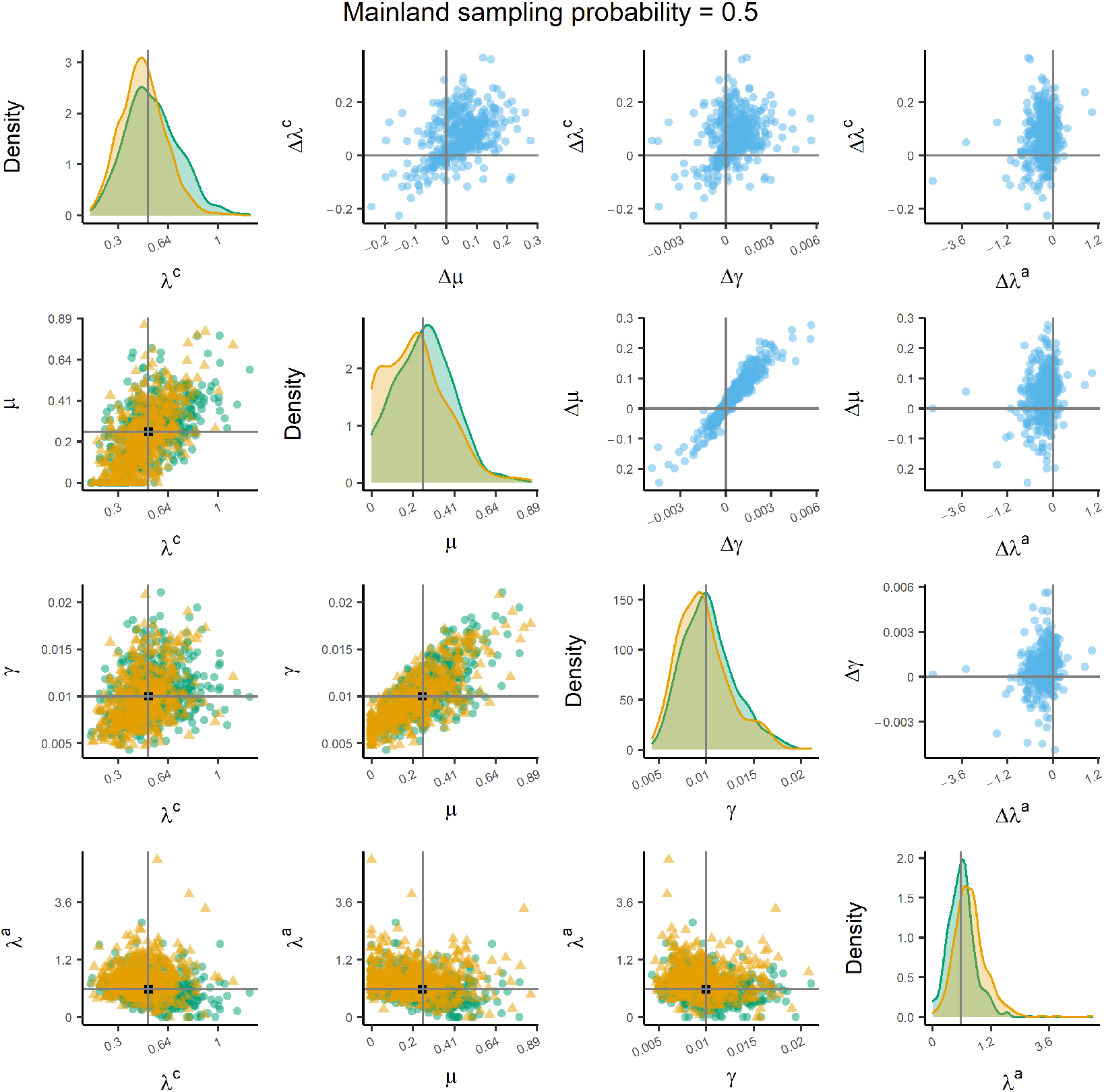
Parameter estimates inferred from the *ideal* and *empirical*_*US*_ data sets of a simulation with a mainland sampling probability of 0.5. The diagonal panels show the distributions of the parameter estimates from the *ideal* (green) and *empirical*_*US*_ (yellow) data sets. The vertical grey line is the true value used to simulate the data. The panels on the lower triangular show the point estimates from the *ideal* (green circles) and *empirical*_*US*_ (yellow triangles) data sets. The black square at the intersection of the grey lines is the true value used to simulate the data. The panels on the upper triangular show the differences between the green circles and yellow triangles (i.e. *ideal* minus *empirical*_*US*_). The grey lines intersect at zero. The *x* -axis on the diagonal panels and both axes on the off-diagonal panels are transformed with inverse hyperbolic sine transformation (ln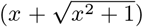) to display data (including zeros and negative values) over several orders of magnitude; the tick labels present the untransformed values. These results were simulated with *K*^*′*^ = 5 and are qualitatively similar to the results from *K*^*′*^ = 50. Anagenesis (*λ*^*a*^) is fixed under certain conditions (see supplementary methods).

**Figure S10:**
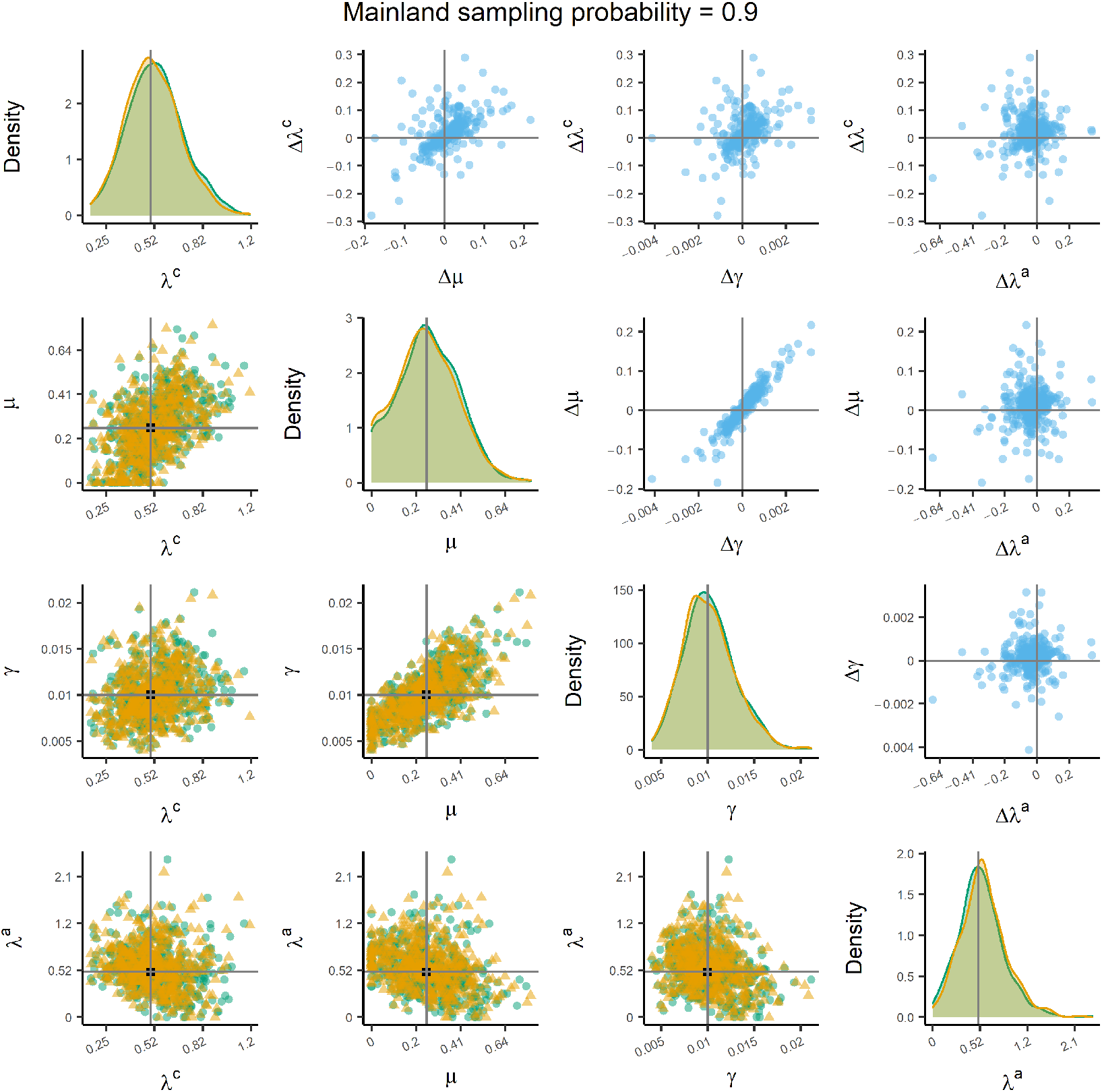
Parameter estimates inferred from the *ideal* and *empirical*_*US*_ data sets of a simulation with a mainland sampling probability of 0.9. The diagonal panels show the distributions of the parameter estimates from the *ideal* (green) and *empirical*_*US*_ (yellow) data sets. The vertical grey line is the true value used to simulate the data. The panels on the lower triangular show the point estimates from the *ideal* (green circles) and *empirical*_*US*_ (yellow triangles) data sets. The black square at the intersection of the grey lines is the true value used to simulate the data. The panels on the upper triangular show the differences between the green circles and yellow triangles (i.e. *ideal* minus *empirical*_*US*_). The grey lines intersect at zero. The *x* -axis on the diagonal panels and both axes on the off-diagonal panels are transformed with inverse hyperbolic sine transformation (ln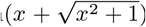) to display data (including zeros and negative values) over several orders of magnitude; the tick labels present the untransformed values. These results were simulated with *K*^*′*^ = 5 and are qualitatively similar to the results from *K*^*′*^ = 50. Anagenesis (*λ*^*a*^) is fixed under certain conditions (see supplementary methods).

**Figure S11:**
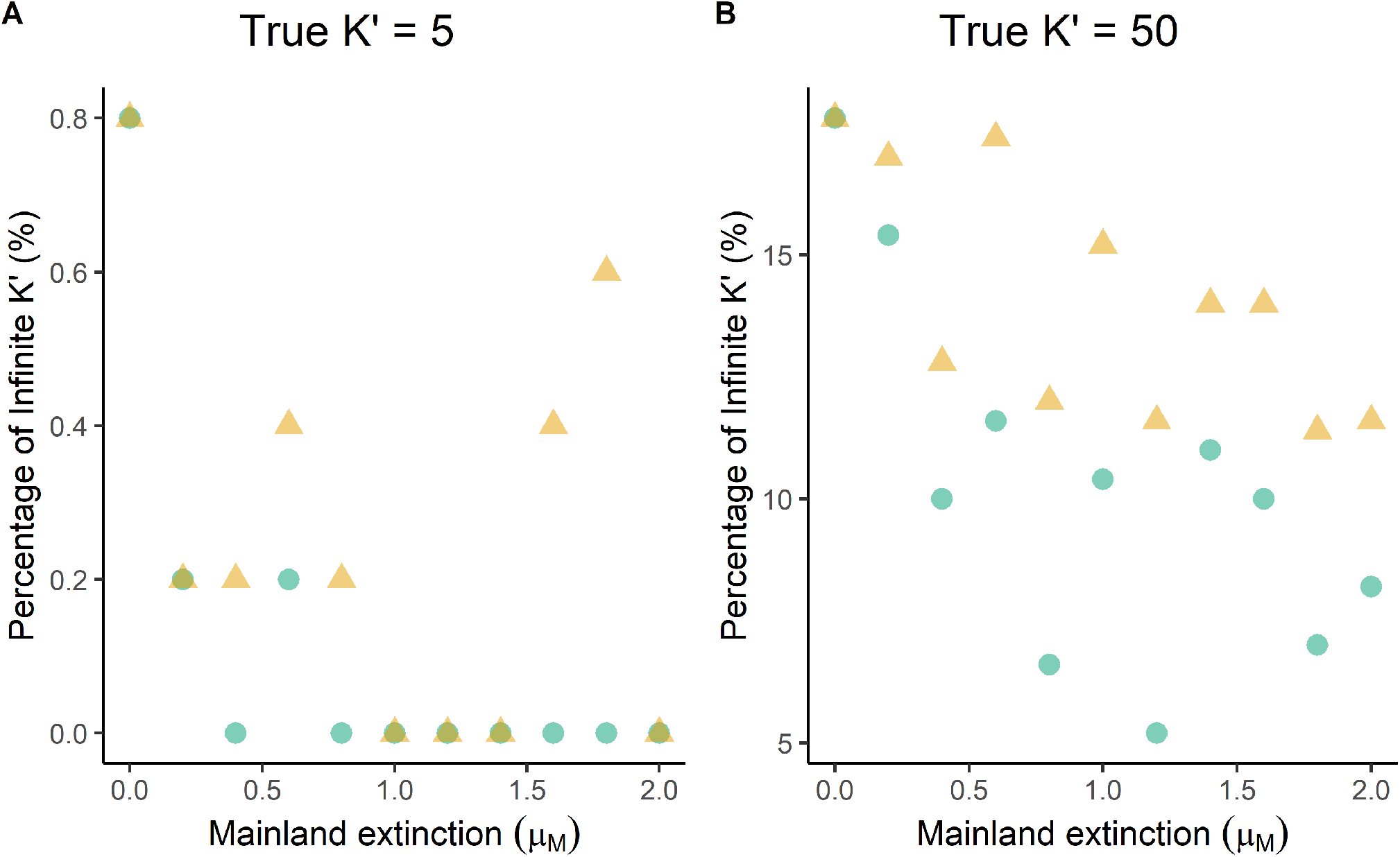
Percentage of carrying capacity estimates that were infinite for both (A) *K*^*′*^ = 5 and (B) *K*^*′*^ = 50, across different rates of mainland extinction (*μ*_*M*_).

**Figure S12:**
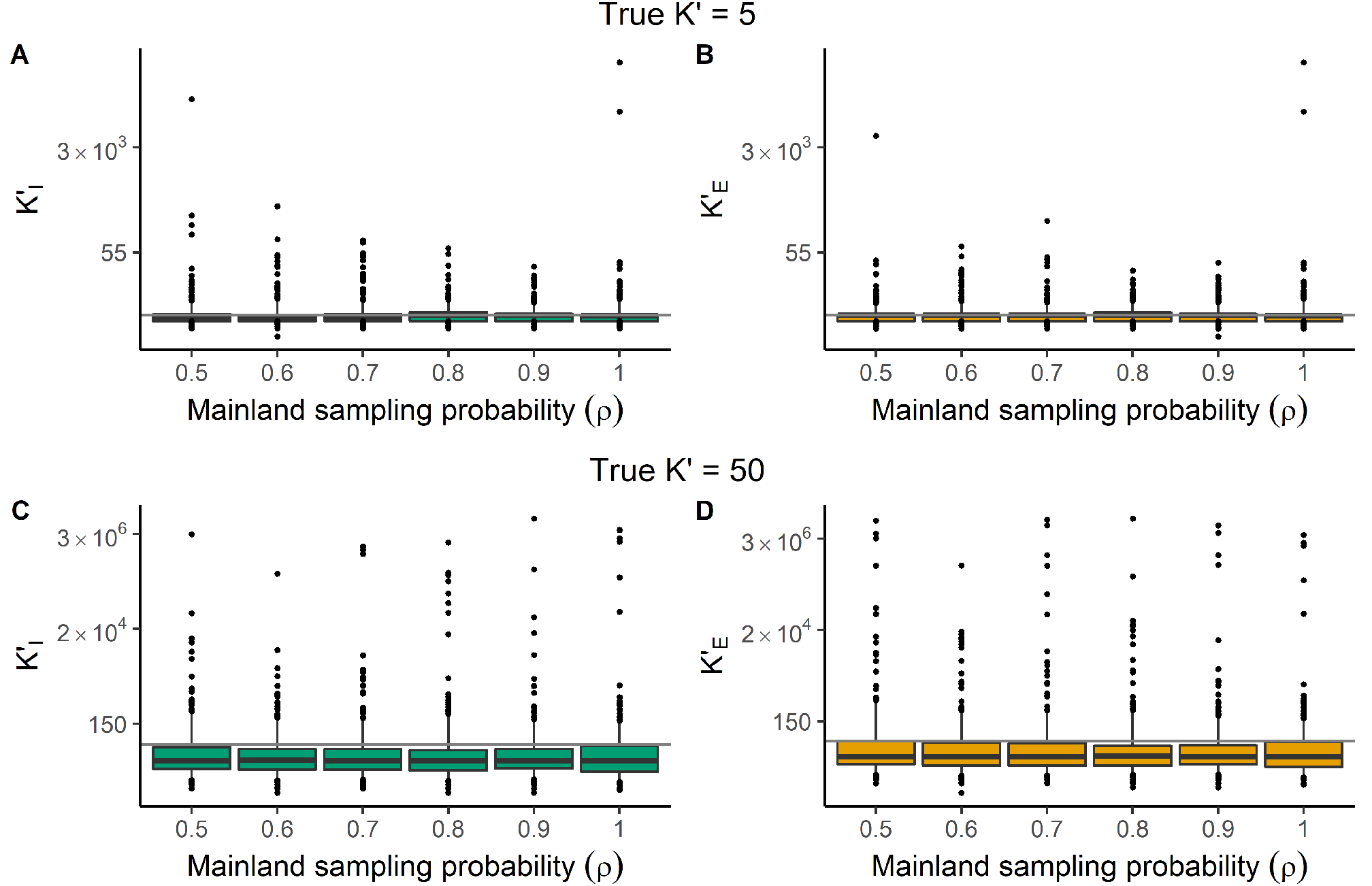
Estimates of carrying capacity parameter (*K*^*′*^) for (A, C) *ideal* data sets (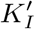, green) and (B, D) *empirical*_*US*_ data sets (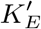, yellow) under two simulated carrying capacities *K*^*′*^ = 5 (A, B) and *K*^*′*^ = 50 (C, D), plotted for different values of mainland sampling probability (*ρ*). The grey horizontal line is the true value. Boxes show the median, 25th and 75th percentiles, while the whiskers extend to the 5th and 95th percentiles. The dots represent points outside of this range. The *y* -axis is plotted on a natural log scale to display data over several orders of magnitude.

**Figure S13:**
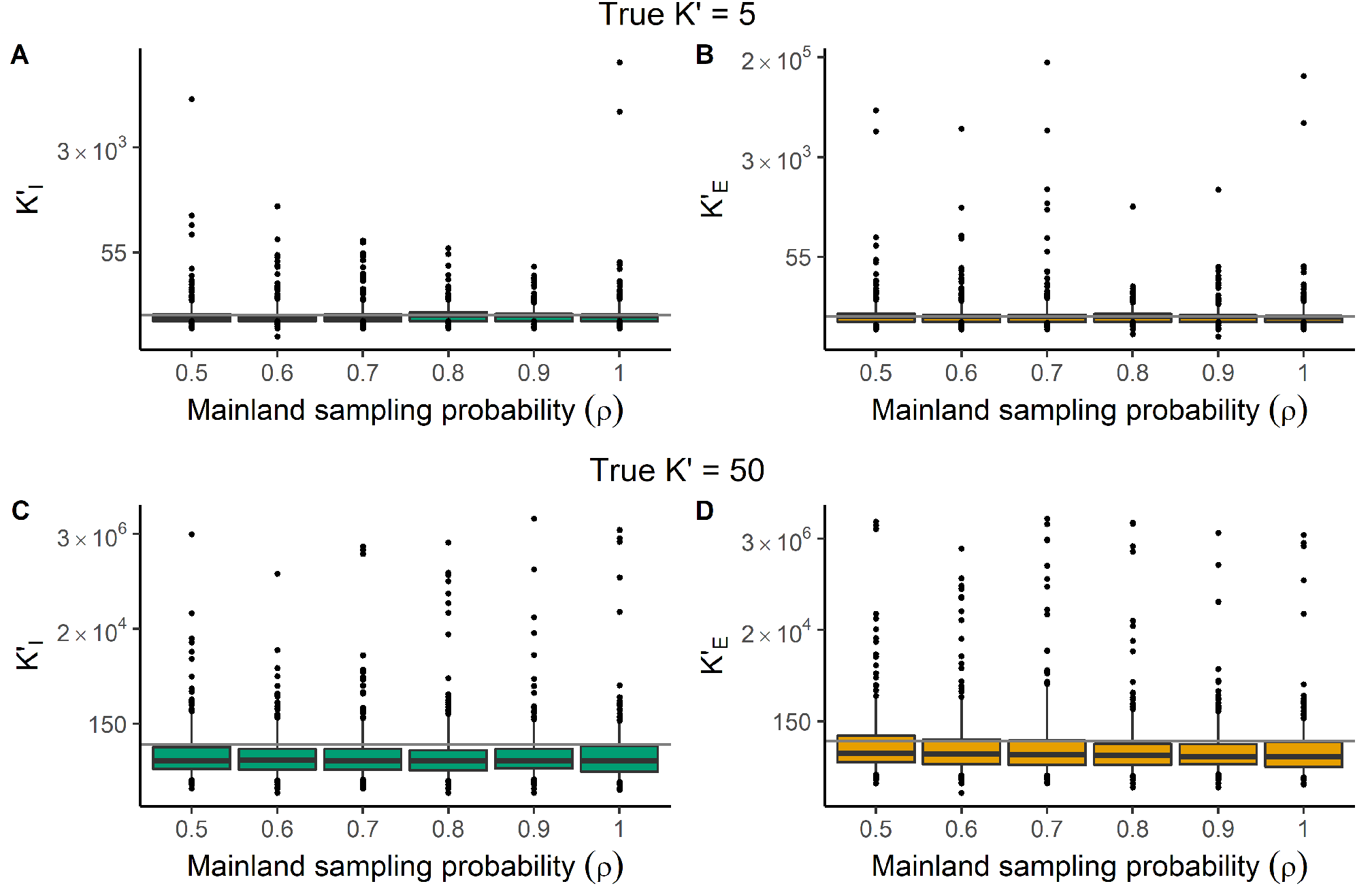
Estimates of carrying capacity parameter (*K*^*′*^) for (A, C) *ideal* data sets (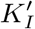, green) and (B, D) *empirical*_*UD*_ data sets (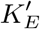, yellow) under two simulated carrying capacities *K*^*′*^ = 5 (A, B) and *K*^*′*^ = 50 (C, D), plotted for different values of mainland sampling probability (*ρ*). The grey horizontal line is the true value. Boxes show the median, 25th and 75th percentiles, while the whiskers extend to the 5th and 95th percentiles. The dots represent points outside of this range. The *y* -axis is plotted on a natural log scale to display data over several orders of magnitude.

## Appendix A

### Ideal vs empirical datasets

In this section the differences between *ideal* and *empirical* datasets are explained. An *ideal* dataset is the complete information of the timing of all events without any error. The *empirical* dataset is the *ideal* data, altered to represent what an empiricist would see when compiling a data set from reconstructed phylogenetic trees (i.e. no extinct species) to apply the DAISIE inference (maximum likelihood) model. In each island-mainland scenario the phylogenetic information of the island and mainland are shown. The assumptions of the DAISIE model are: there is no back-colonisation from the island to the mainland; cladogenetic speciation on the island or on the mainland produces two new species and the ancestor is no longer present (i.e. symmetric speciation); there is no divergence in the mainland species (i.e. genetically homogeneous) before they immigrate to the island; there is no incomplete lineage sorting so all species are correctly delineated in the tree; the timing of immigration events can be detected as the founder effect and evolution in isolation would leave a genetic signature. For the last assumption, the empirical data assumes that the phylogeny is complete at the population level, in order to determine divergence of populations of the same species on the island and mainland.

The DAISIE inference model needs information on the status of the clade (stac) in the following way:

stac 0: Empty island

stac 1: Non-endemic with unknown colonisation time but with a maximum to this colonisation time

stac 2: Endemic singleton or endemic clade

stac 3: Endemic singleton or endemic clade with one or more re-colonisations of the same mainland species

stac 4: Non-endemic singleton with known colonisation time

stac 5: Endemic singleton with unknown colonisation time, but with a maximum to this colonisation time

stac 6: Endemic clade with unknown colonisation time, but with a maximum to this colonisation time

stac 7: Endemic singleton or endemic clade and one or more re-colonisations of the same mainland species with unknown colonisation times, but with a maximum to this colonisation time

stac 8: Non-endemic with unknown colonisation time, but with a maximum and minimum to this colonisation time

stac 9: Endemic singleton with unknown colonisation time, but with a maximum and minimum to this colonisation time

In the *ideal* data only stac 2, 3 and 4 are assigned because it is known when the species colonises the island. However, in the *empirical* data stac 1, 5 and 6 can also be assigned when the colonisation time is not known but only a maximum (often island age). This is explained below.

### Empty Island

If the island is empty at the end of the simulation, either through the failure of any mainland species to colonise, or if all the colonising species went extinct before the present then the *empirical* data is the same as the *ideal* data, which is that all the empiricist knows is that at the present there are no species. *Empirical* and *Ideal* data are assigned stac 0.

**Figure A1:**
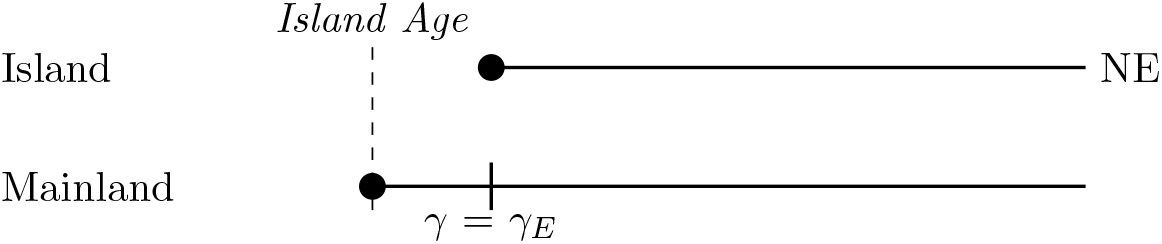
If the mainland species immigrates and does not go extinct or speciate, and no events happen on the island, the island species is non-endemic (NE), and the colonisation time in the *empirical* data (*γ*_*E*_) is the same as the colonisation time in the *ideal* data (*γ*). *Empirical* and *ideal* data are assigned stac 4.

**Figure A2:**
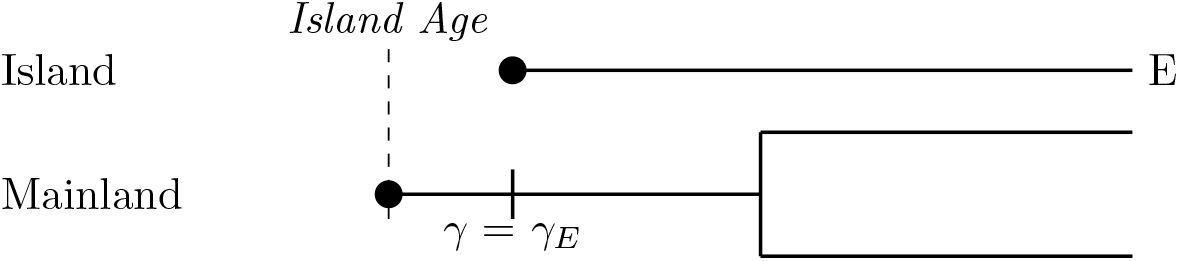
If the mainland species immigrates to the island then undergoes cladogenesis on the mainland and the descendent species do not go extinct and no events happen on the island, the island species is endemic (E) and in the *empirical* data the colonisation time (*γ*_*E*_) is the same as the colonisation time in the *ideal* data (*γ*). *Empirical* and *ideal* data are assigned stac 2.

**Figure A3:**
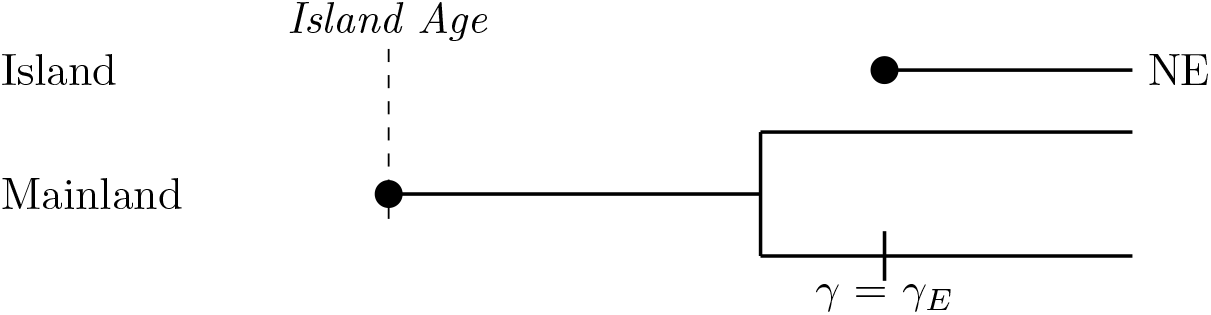
If the mainland species undergoes speciation and then one of the descendent species immigrates to the island and both the descendent species do not go extinct and no events happen on the island, the island species is non-endemic (NE) and in the *empirical* data the colonisation time (*γ*_*E*_) is the same as the colonisation time in the *ideal* data (*γ*). *Empirical* and *ideal* data are assigned stac 4.

**Figure A4:**
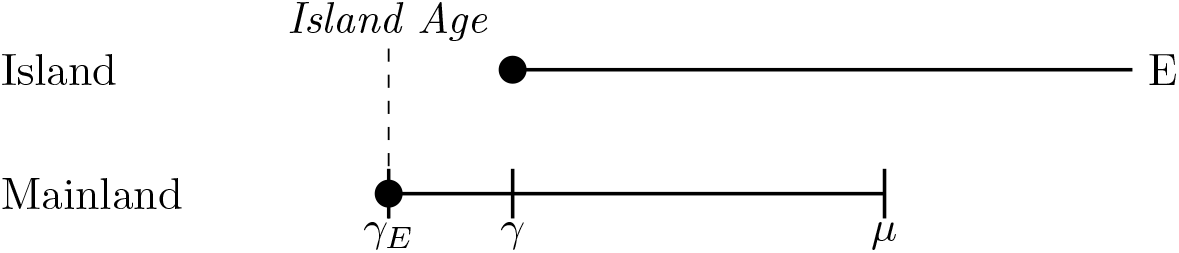
If the mainland species colonises the island and then goes extinct on the mainland (*μ*), without having speciated, and no events happen on the island, the island species is endemic (E) and in the *empirical* data the colonisation time (*γ*_*E*_) is the maximum age of the island. *Ideal* data is assigned stac 2 and *empirical* data is assigned stac 5.

**Figure A5:**
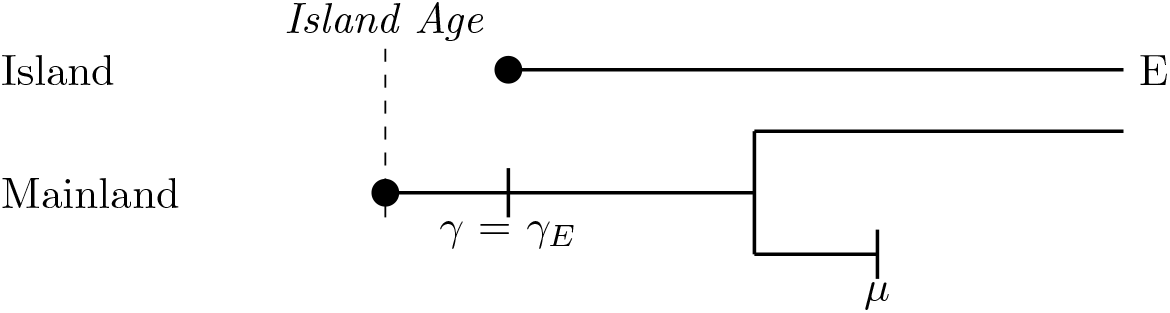
If the mainland species colonises the island and then undergoes speciation and one of the descendant species goes extinct (*μ*), and no events happen on the island, the island species is endemic (E) and in the *empirical* data the colonisation time (*γ*_*E*_) is the same as the colonisation time in the *ideal* data (*γ*). *Ideal* and *empirical* data are assigned stac 2.

**Figure A6:**
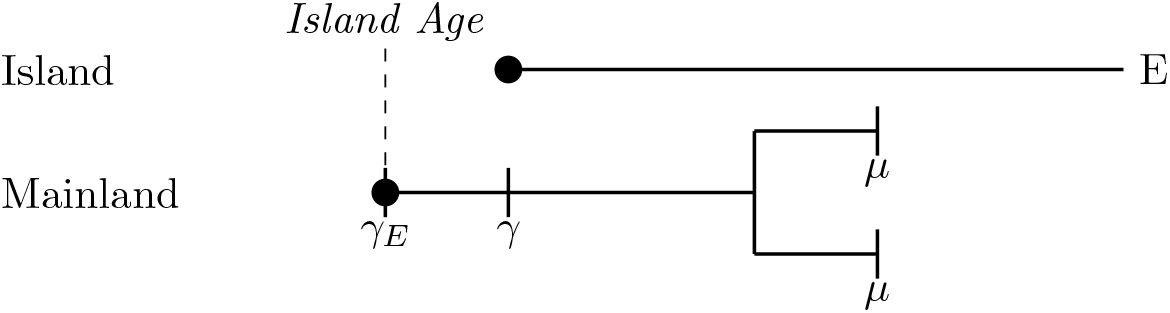
If the mainland species colonises the island and then undergoes speciation and both of the descendant species goes extinct (*μ*), and no events happen on the island, the island species is endemic (E) and in the *empirical* data the colonisation time (*γ*_*E*_) is the maximum age of the island. *Ideal* data is assigned stac 2 and *empirical* data is assigned stac 5.

**Figure A7:**
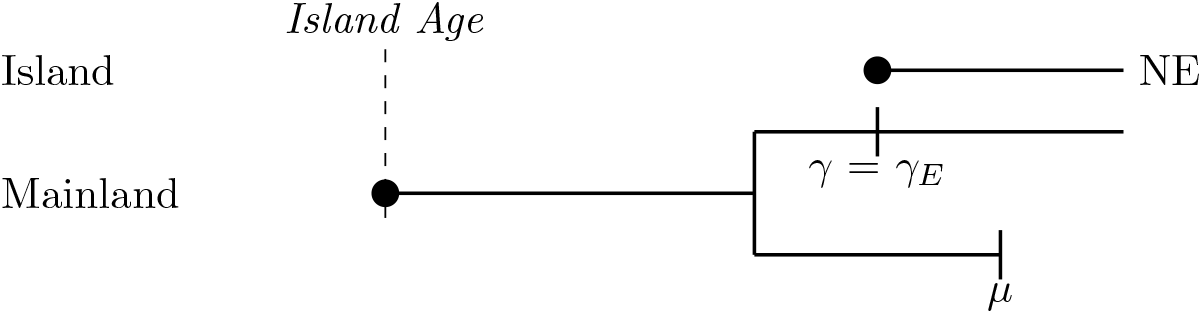
If the mainland species undergoes speciation and then one of the descendant species immigrates to the island and the other descendant goes extinct (*μ*) and no events happen on the island, the island species is non-endemic (NE) and in the *empirical* data the colonisation time (*γ*) is the same as the colonisation time in the *ideal* data (*γ*). *Ideal* and *empirical* data are assigned stac 4.

**Figure A8:**
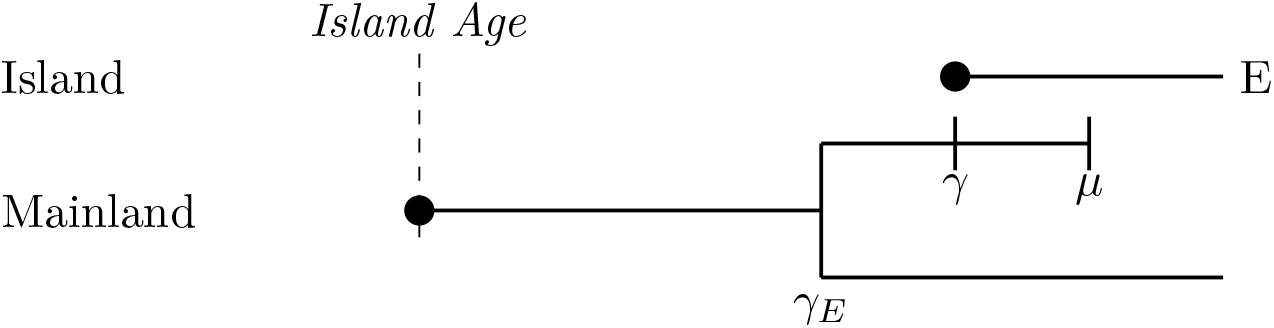
If the mainland species undergoes speciation and then one of the descendant species immigrates to the island and then goes extinct on the mainland (*μ*), and the other descendent survives, and no events happen on the island, the island species is endemic (E) and in the *empirical* data the colonisation time (*γ*_*E*_) is the branching time on the mainland. *Ideal* and *empirical* data are assigned stac 2.

### Single species on the island with no island events

**Figure A9:**
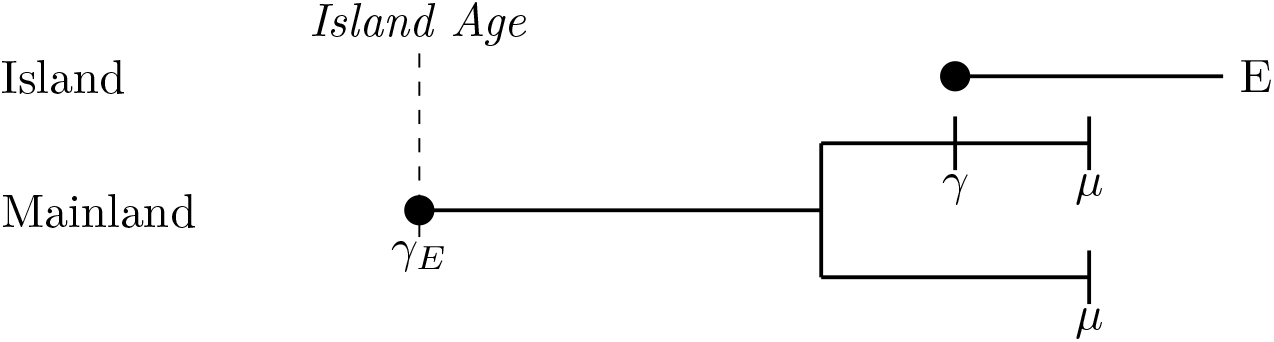
If the mainland species undergoes speciation and then one of the descendant species immigrates to the island and then both descendants go extinct (*μ*), and no events happen on the island, the island species is endemic (E) and in the *empirical* data the colonisation time (*γ*_*E*_) is the maximum age of the island. *Ideal* data is assigned stac 2 and *empirical* data is assigned stac 5.

### Single species on the island with anagenesis

**Figure A10:**
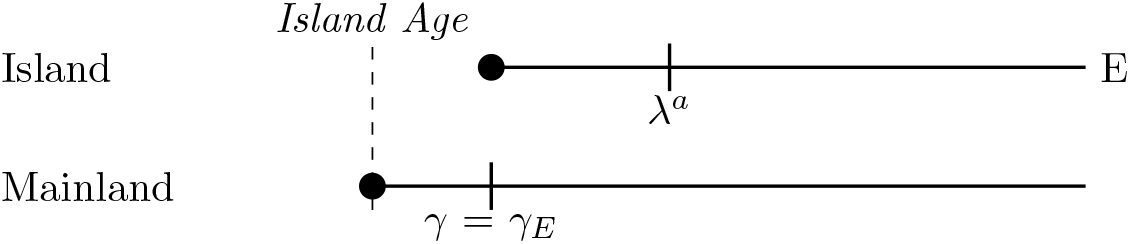
If the mainland species immigrates to the island and does not go extinct or speciate, and the island species undergoes anagenesis, the island species is endemic (E), and in the *empirical* data the colonisation time (*γ*_*E*_) is the same as the colonisation time in the *ideal* data (*γ*). *Ideal* and *empirical* data is assigned stac 2.

**Figure A11:**
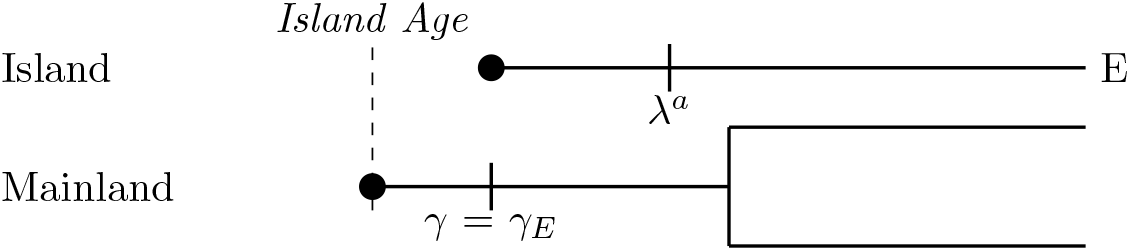
If the mainland species immigrates to the mainland and then undergoes speciation, with both mainland descendants surviving to the present, and the island species undergoes anagenesis, the island species is endemic (E) and in the *empirical* data the colonisation time (*γ*_*E*_) is the same as the colonisation time in the *ideal* data (*γ*). *Ideal* and *empirical* data are assigned stac 2.

**Figure A12:**
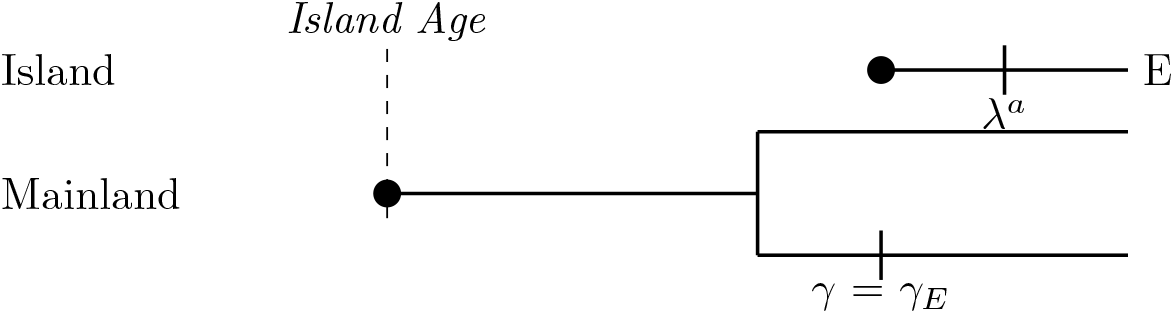
If the mainland species undergoes speciation and then one of the descendent species immigrates to the island and both the descendent species do not go extinct and the island species undergoes anagenesis, the island species is endemic (E) and in the *empirical* data the colonisation time (*γ*_*E*_) is the same as the colonisation time in the *ideal* data (*γ*). *Ideal* and *empirical* data are assigned stac 2.

**Figure A13:**
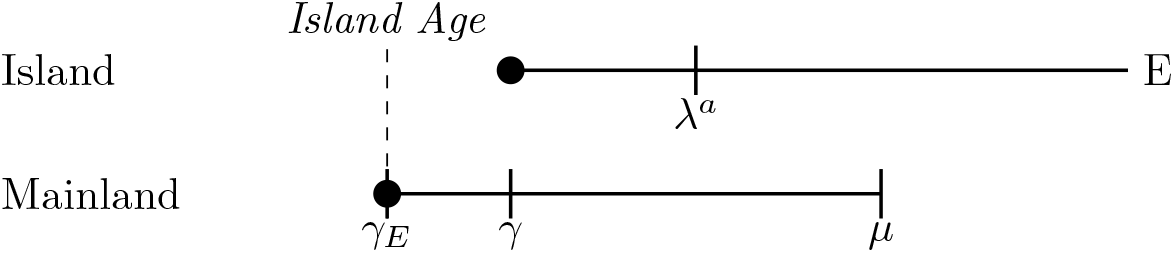
If the mainland species immigrates to the island (*γ*) and then goes extinct (*μ*) on the mainland, and the island species undergoes anagenesis, the island species is endemic (E), and in the *empirical* data the colonisation time (*γ*_*E*_) is the maximum age of the island. *Ideal* data is assigned stac 2 and *empirical* data is assigned stac 5.

**Figure A14:**
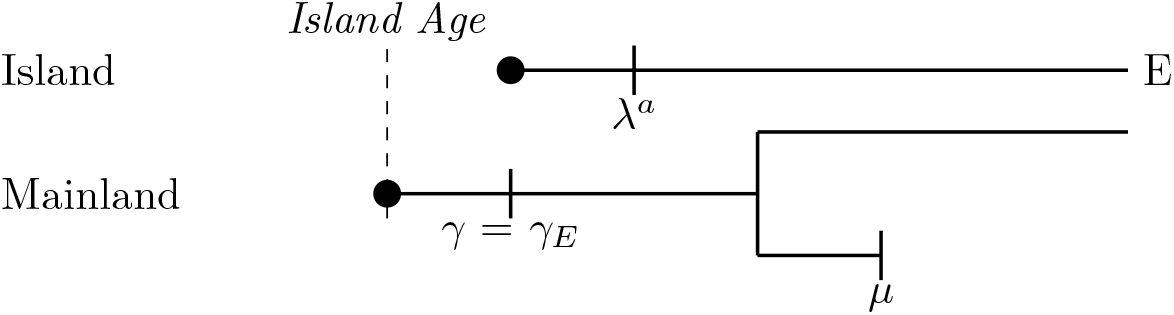
If the mainland species colonises the island (*γ*) and then undergoes speciation, and one or both of the descendent species go extinct on the mainland (*μ*), and the island species undergoes anagenesis, the island species is endemic (E), and in the *empirical* data the colonisation time (*γ*_*E*_) is the same as the colonisation time in the *ideal* data (*γ*). *Ideal* and *empirical* is assigned stac 2.

**Figure A15:**
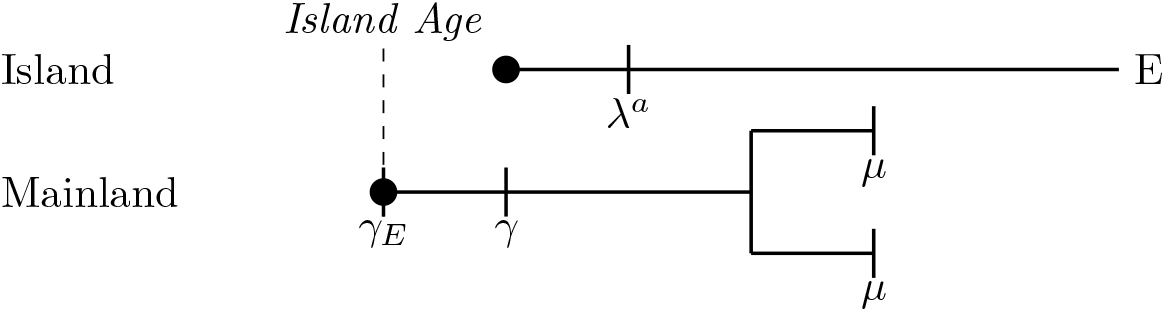
If the mainland species colonises the island and then undergoes speciation, and one or both of the descendent species go extinct (*μ*), and the island species undergoes anagenesis, the island species is endemic (E), and in the *empirical* data the colonisation time (*γ*_*E*_) is the maximum age of the island. *Ideal* data is assigned stac 2 and *empirical* data is assigned stac 5.

**Figure A16:**
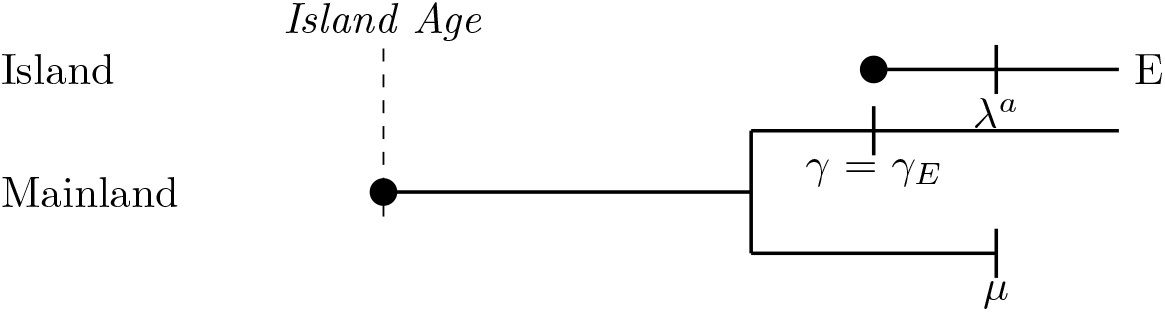
If the mainland species undergoes speciation and then one of the descendant species immigrates to the island and the other descendant goes extinct (*μ*) and the island species undergoes anagenesis, the island species is endemic (E) and in the *empirical* data the colonisation time (*γ*_*E*_) is the same as the colonisation time in the *ideal* data (*γ*). *Ideal* and *empirical* data are assigned stac 2.

**Figure A17:**
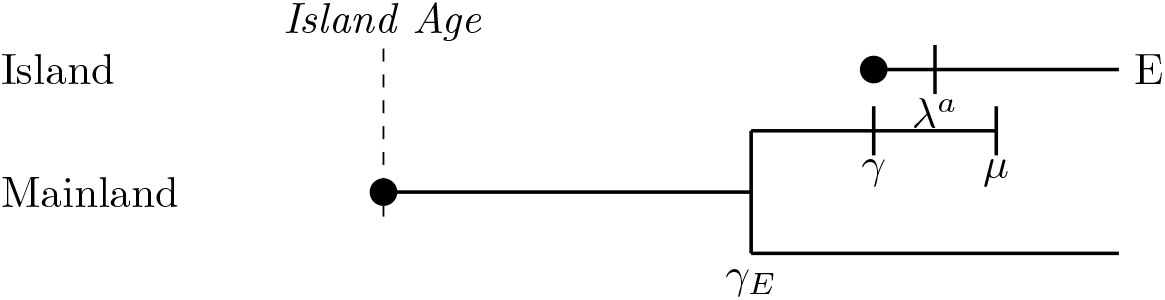
If the mainland species undergoes speciation and then one of the descendant species immigrates to the island and then goes extinct on the mainland (*μ*), and the other descendent survives, and the island species undergoes anagenesis, the island species is endemic (E) and in the *empirical* data the colonisation time (*γ*_*E*_) is the branching time on the mainland. *Ideal* and *empirical* are assigned stac 2.

**Figure A18:**
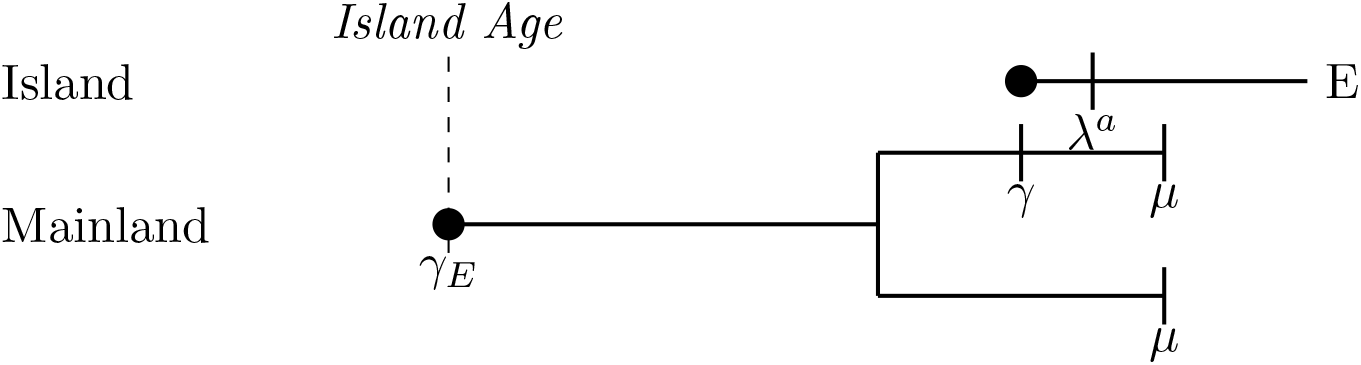
If the mainland species undergoes speciation and then one of the descendant species immigrates to the island (*γ*) and then both descendants go extinct on the mainland (*μ*), and the island species undergoes anagenesis, the island species is endemic (E) and in the *empirical* data the colonisation time (*γ*_*E*_) is the maximum age of the island. *Ideal* data is assigned stac 2 and *empirical* data is assigned stac 5.

### Clade on the island

**Figure A19:**
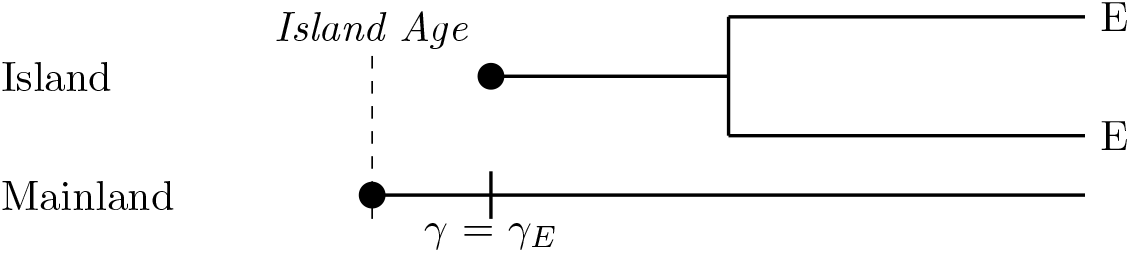
If the mainland species colonises the island and does not go extinct or speciate, and the island species undergoes cladogenesis the species are endemic (E), and in the *empirical* data the colonisation time (*γ*_*E*_) is the same as the colonisation time in the *ideal* data (*γ*). *Ideal* and *empirical* data are assigned stac 2.

**Figure A20:**
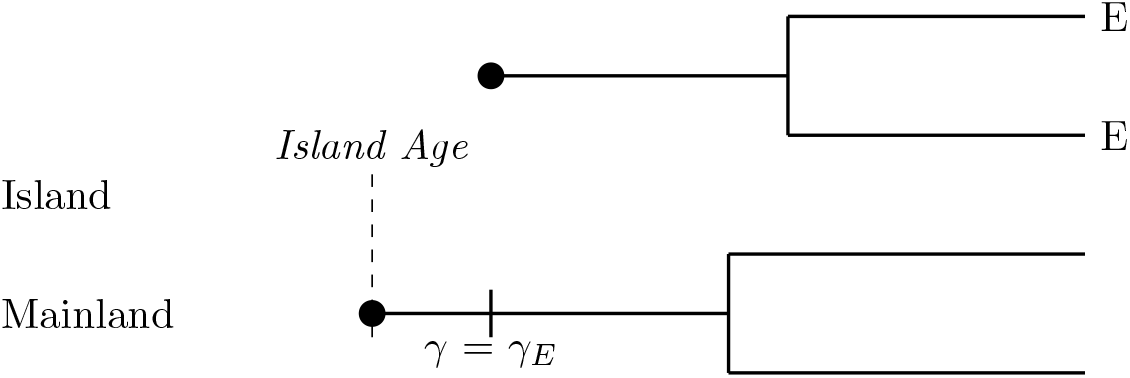
If the mainland species colonises the island and then undergoes speciation and neither of the descendent species go extinct and the species on the island undergoes cladogenesis both species are endemic (E) and in the *empirical* data the colonisation time (*γ*_*E*_) is the same as the colonisation time in the *ideal* data (*γ*). *Ideal* and *empirical* data are assigned stac 2.

**Figure A21:**
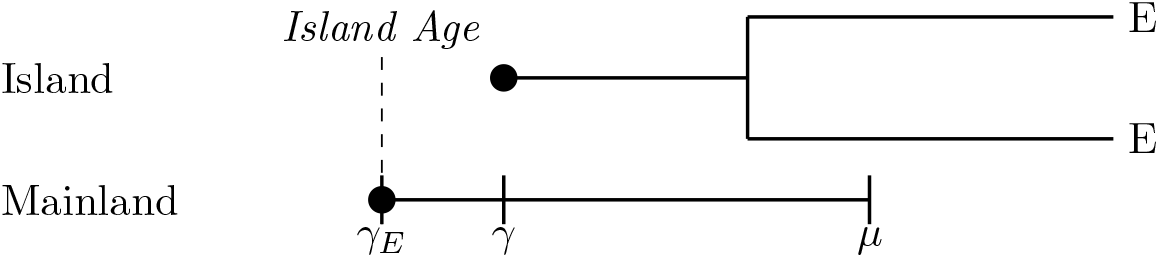
If the mainland species colonises the island (*γ*) and then goes extinct on the mainland (*μ*), and the island species undergoes cladogenesis the species are endemic (E), and in the *empirical* data the colonisation time (*γ*_*E*_) is the maximum age of the island. *Ideal* data is assigned stac 2 and *empirical* data is assigned stac 6.

**Figure A22:**
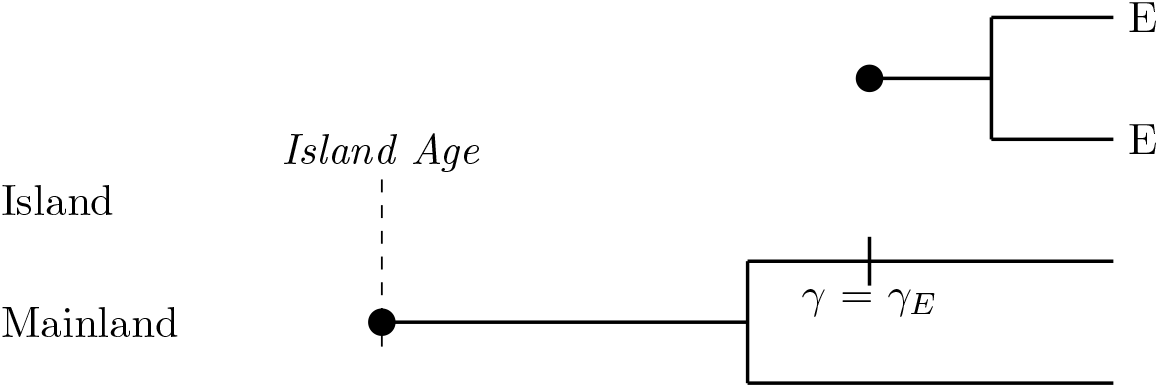
If the mainland species undergoes speciation and then one of the descendant species colonises the island and both descendent species survive to the present and the island species undergoes cladogenesis, the two island species are endemic (E), and in the *empirical* data the colonisation time (*γ*_*E*_) is the same as the colonisation time in the *ideal* data (*γ*). *Ideal* and *empirical* data are assigned stac 2.

**Figure A23:**
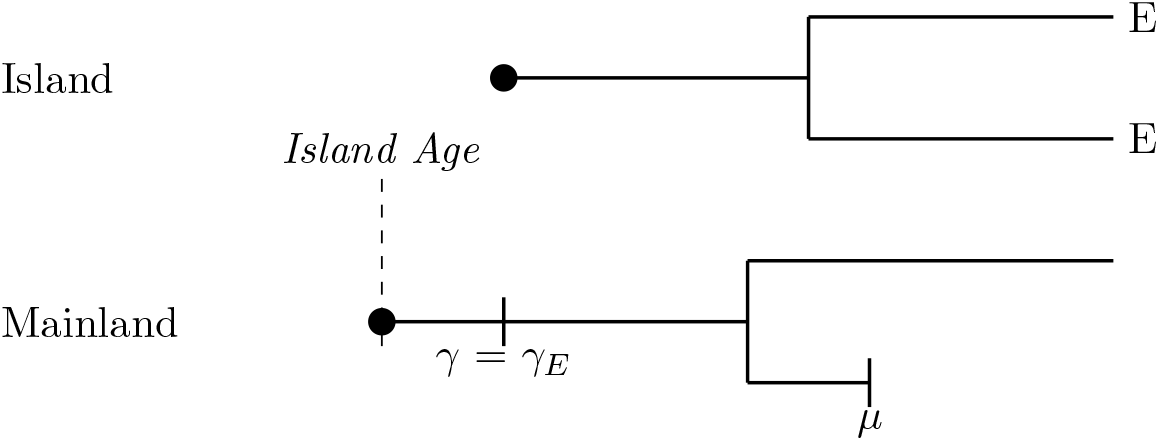
If the mainland species colonises the island, then undergoes speciation, and one of the descendent species goes extinct (*μ*) and the island species undergoes cladogenesis on the island, the two island species are endemic (E), and in the *empirical* data the colonisation time (*γ*_*E*_) is the same as the colonisation time in the *ideal* data (*γ*). *Ideal* and *empirical* data are assigned stac 2.

**Figure A24:**
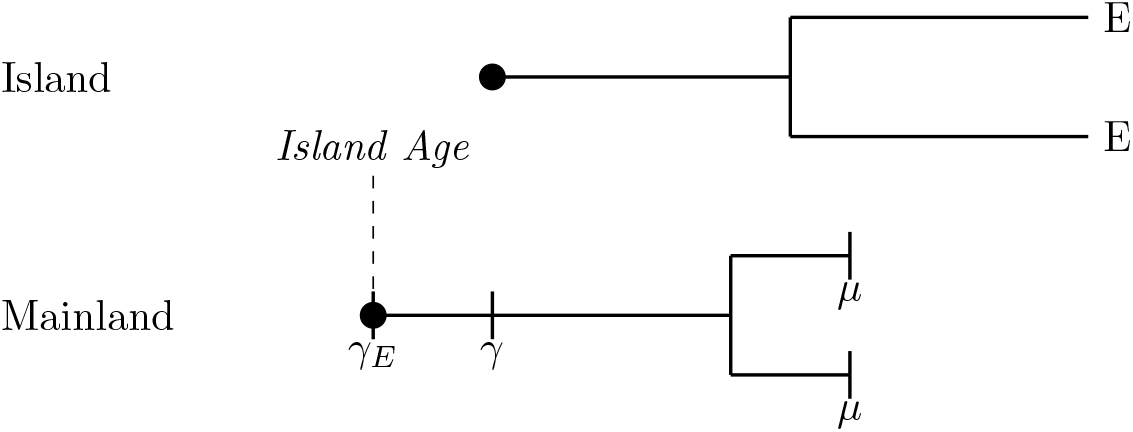
If the mainland species colonises the island (*γ*), then undergoes speciation, and both of the descendent species go extinct on the mainland (*μ*) and the island species undergoes cladogenesis on the island, the two island species are endemic (E), and in the *empirical* data the colonisation time (*γ*_*E*_) is the maximum age of the island. *Ideal* data is assigned stac 2 and *empirical* data is assigned stac 6.

**Figure A25:**
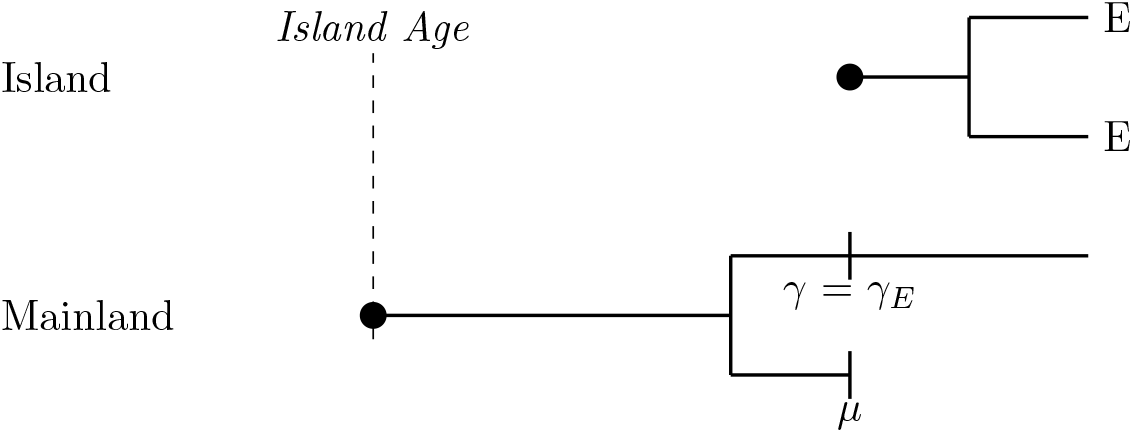
If the mainland species undergoes speciation, then colonises the island, and one of the descendent species goes extinct on the mainland (*μ*) and the island species undergoes cladogenesis on the island, the two island species are endemic (E), and in the *empirical* data the colonisation time (*γ*_*E*_) is the same as the colonisation time in the *ideal* data (*γ*). *Ideal* and *empirical* data are assigned stac 2

**Figure A26:**
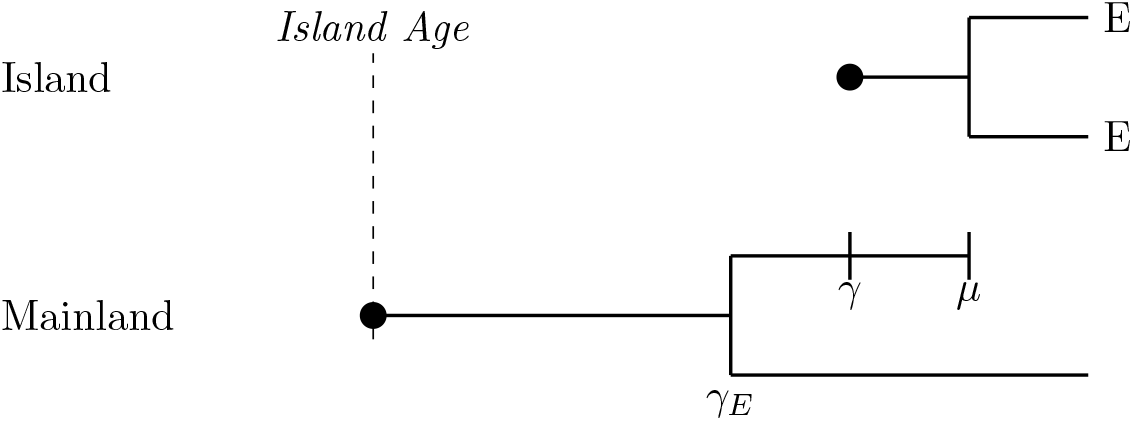
If the mainland species undergoes speciation, then one descendant colonises the island (*γ*), and then goes extinct on the mainland (*μ*) and the other descendant survives to the present, and the island species undergoes cladogenesis on the island, the two island species are endemic (E), and in the *empirical* data (*γ*_*E*_) the colonisation time is the branching time on the mainland. *Ideal* and *empirical* data are assigned stac 2.

**Figure A27:**
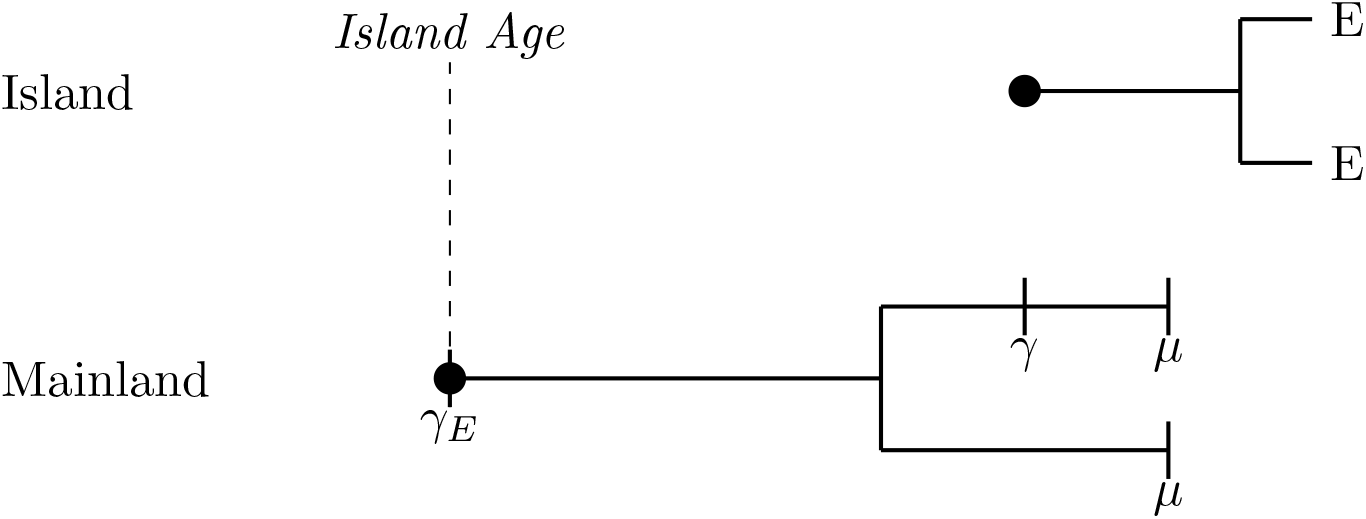
If the mainland species undergoes speciation, then colonises the island (*γ*), and both of the descendent species go extinct on the mainland (*μ*) and the island species undergoes cladogenesis on the island, the two island species are endemic (E), and in the *empirical* data the colonisation time (*γ*_*E*_) is the maximum age of the island. *Ideal* data is assigned stac 2 and *empirical* data is assigned stac 6.

### Colonisation of the same mainland species after cladogenesis or anagenesis on the island

**Figure A28:**
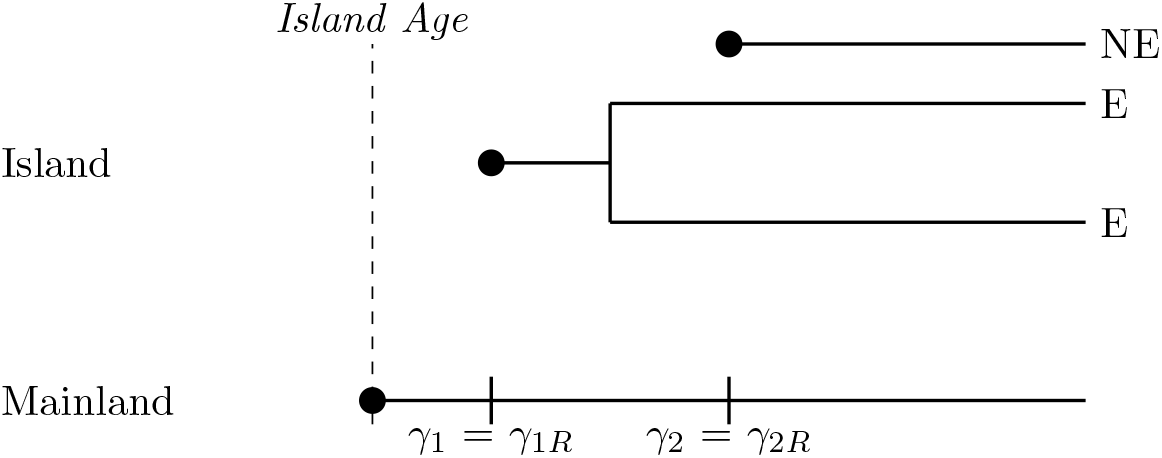
If the mainland species does not speciate or go extinct, and the mainland species colonises the island twice, the second colonisation taking place after the island species has undergone cladogenesis, the two species in the island clade are considered endemic (E), and the second immigrant is non-endemic (NE). The colonisation times in the *empirical* data (*γ*_1*E*_) for the island clade is is the same as the colonisation time in the *ideal* data (*γ*_1_) and for the non-endemic singleton the colonisation time in the *empirical* data (*γ*_2*E*_) is the same as the colonisation time in the *ideal* data (*γ*_2_). *Ideal* and *empirical* data are assigned stac 3.

**Figure A29:**
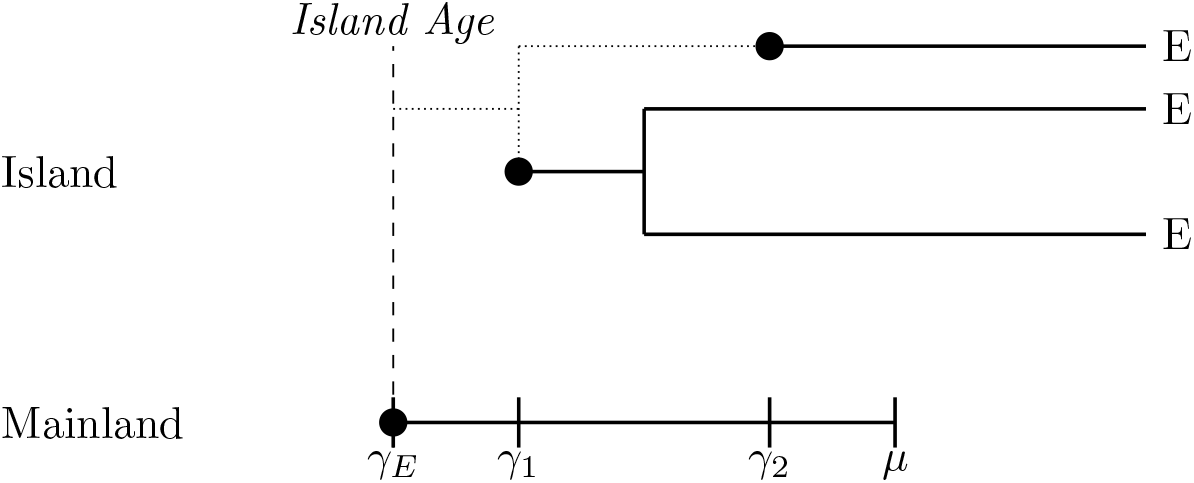
If the mainland species colonises the island twice and then goes extinct (*μ*) on the mainland, and between the immigration events the first island species underwent cladogenesis, all species on the island are considered endemic (E) and in the *empirical* data all species would be considered to arise from a single colonisation time (*γ*_*E*_) at the maximum age of the island. Therefore the second colonisation time (*γ*_2_) is lost, both colonists form a single clade (dotted line), and the singleton endemic is assumed to have colonised anywhere from the first immigration event. *Ideal* data is assigned stac 3 and *empirical* data is assigned stac 6.

**Figure A30:**
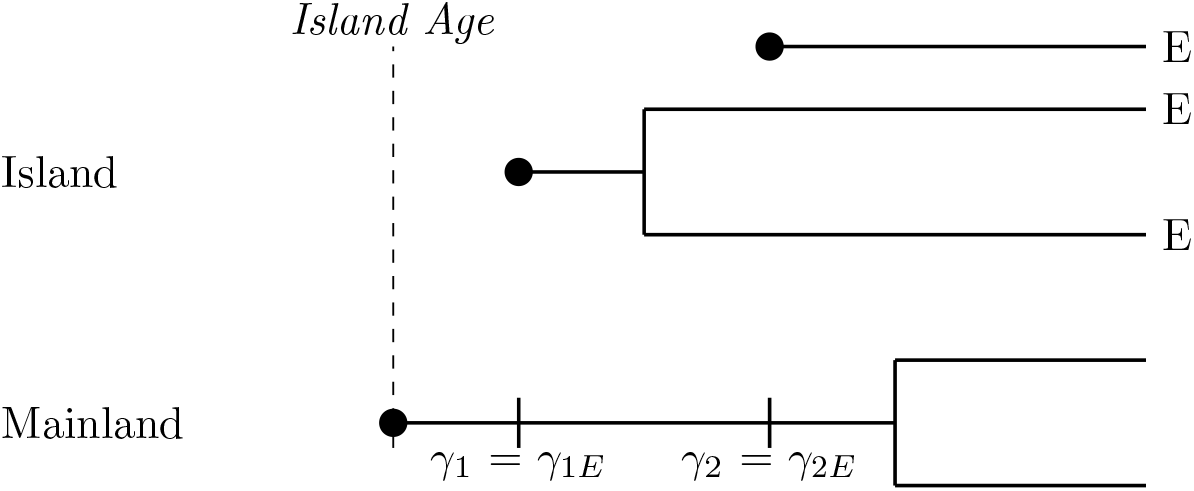
If the mainland species immigrates to the island twice and then undergoes a speciation event, with the island species having undergone cladogenesis between the immigration events, all species on the island are considered endemic (E), and in the *empirical* data the colonisation time (*γ*_1*E*_) for the island clade is the same as the colonisation time in the *ideal* data (*γ*_1_), and for the endemic singleton the *empirical* colonisation time (*γ*_2*E*_) is the same as the colonisation time in the *ideal* data (*γ*_2_), but these are not used in inference because *ideal* and *empirical* data are assigned stac 3.

**Figure A31:**
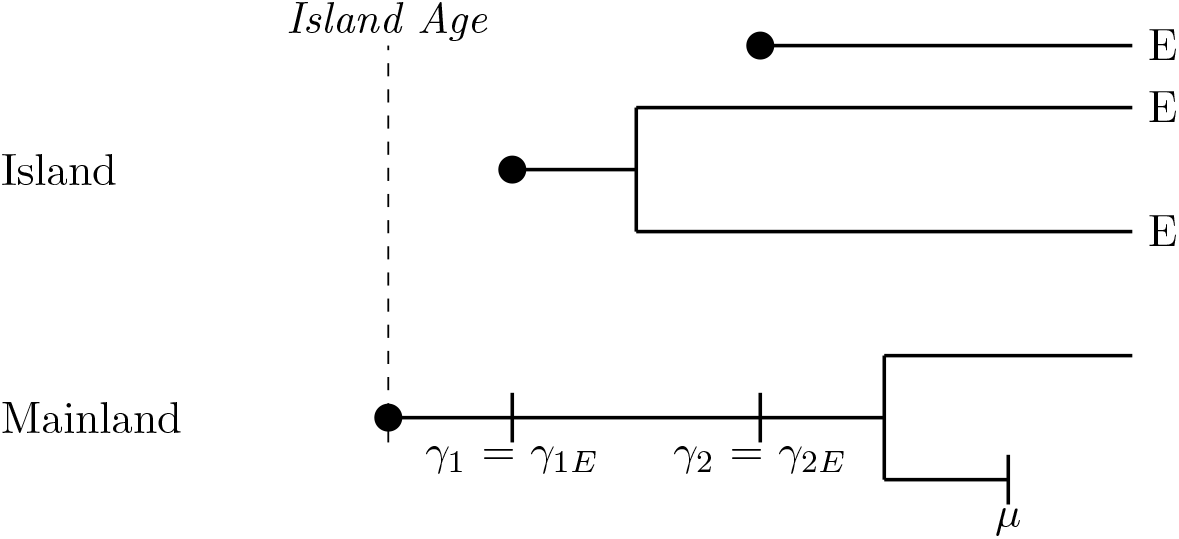
If the mainland species colonises the island twice and then speciates on the mainland, with one of the descendants going extinct (*μ*), and the first island species undergoing cladogenesis before the second immigration event, all species on the island are considered endemic (E), the colonisation times in the *empirical* data (*γ*_1*E*_) for the island clade is the same as the colonisation time in the *ideal* data (*γ*_1_) and for the endemic singleton the colonisation time in the *empirical* data (*γ*_2*E*_) is the same as the colonisation time is the *ideal* data (*γ*_2_), but these are not used in inference, because *ideal* and *empirical* data are assigned stac 3.

**Figure A32:**
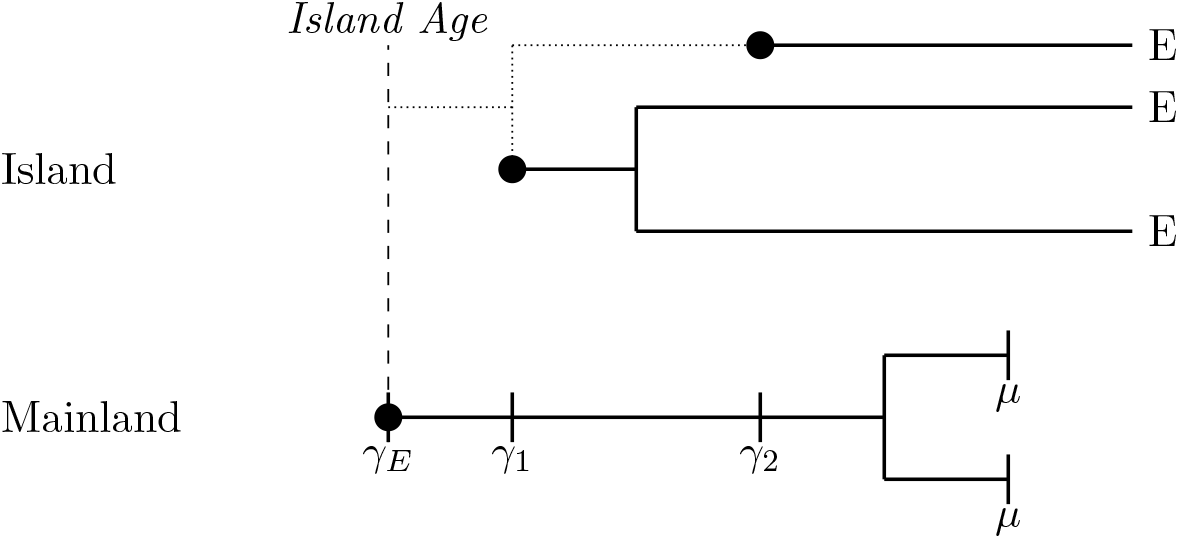
If the mainland species colonises the island twice and then speciates on the mainland, with both of the descendants going extinct (*μ*), and the first island species undergoing cladogenesis before the second immigration event, all species on the island are considered endemic (E) and in the *empirical* data all the species would be considered to arise from a single colonisation time (*γ*_*E*_) at the maximum age of the island. Therefore the second colonisation time (*γ*_2_) is lost and the singleton endemic is assumed to have colonised anywhere since the first immigration event (*γ*_1_). *Ideal* data is assigned stac 3 and *empirical* data is assigned stac 6.

**Figure A33:**
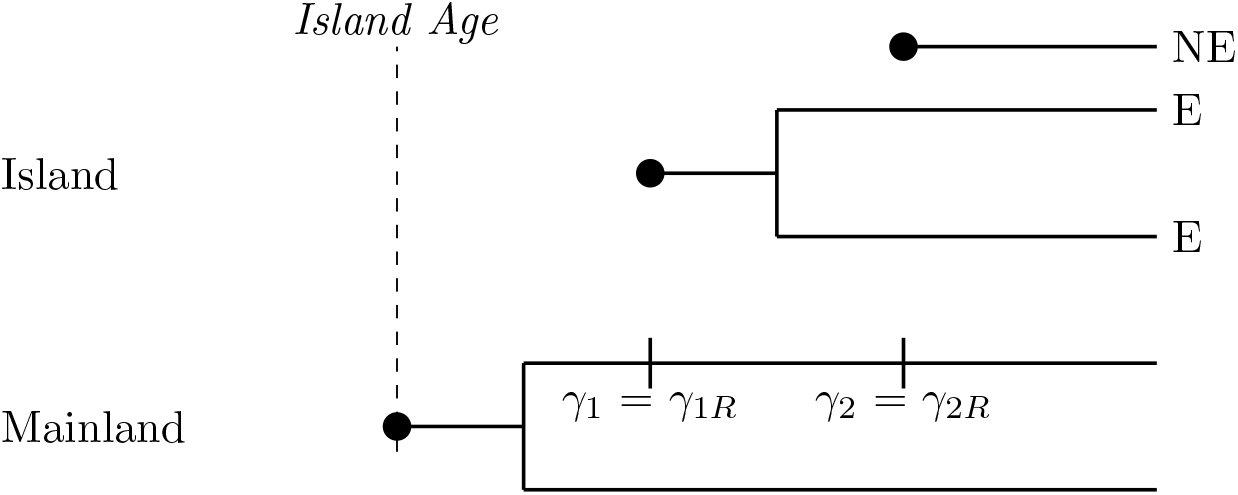
If the mainland species speciates and then one of the descendants colonises the island twice, with both of the descendants surviving, and the first island species undergoing cladogenesis before the second immigration event, the first colonist is an endemic clade (E) and the second colonist is a non-endemic singleton (NE) and the *empirical* is the same as the *ideal. Ideal* and *empirical* data are assigned stac 3, so the second colonisation time is lost with the first setting a maximum to the second.

**Figure A34:**
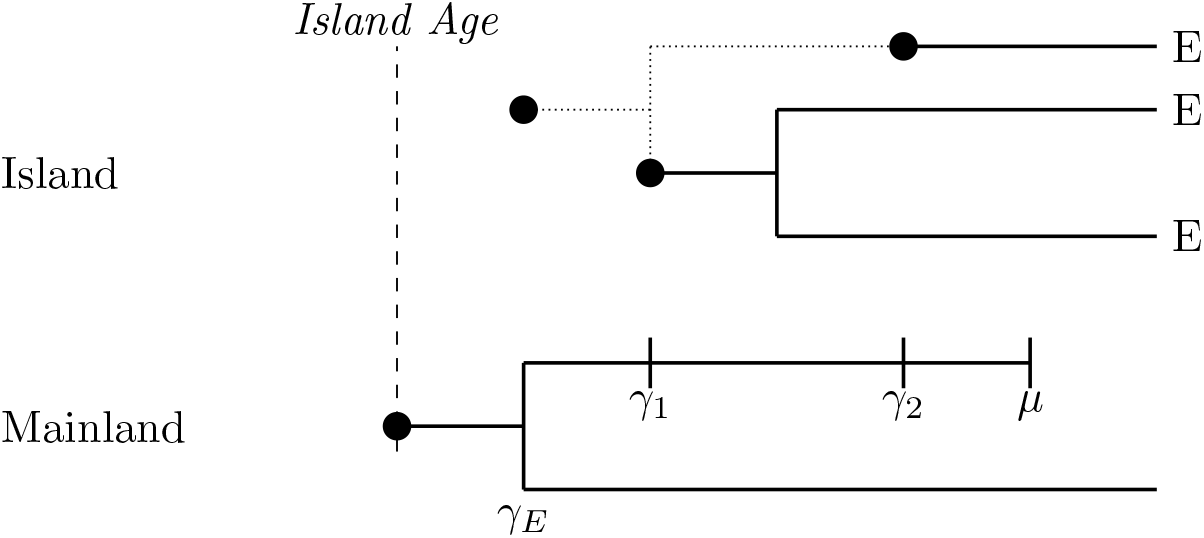
If the mainland species speciates and then one of the descendants colonises the island twice and then goes extinct (*μ*), and the other descendant survives, and the first island species undergoes cladogenesis before the second immigration event, both colonists are endemic (E). In the *empirical* case they are thought to form a single clade with a colonisation time at the branching time on the mainland (*γ*_*E*_). The second colonisation time is lost. *Ideal* data is assigned stac 3 and *empirical* data are assigned stac 2.

**Figure A35:**
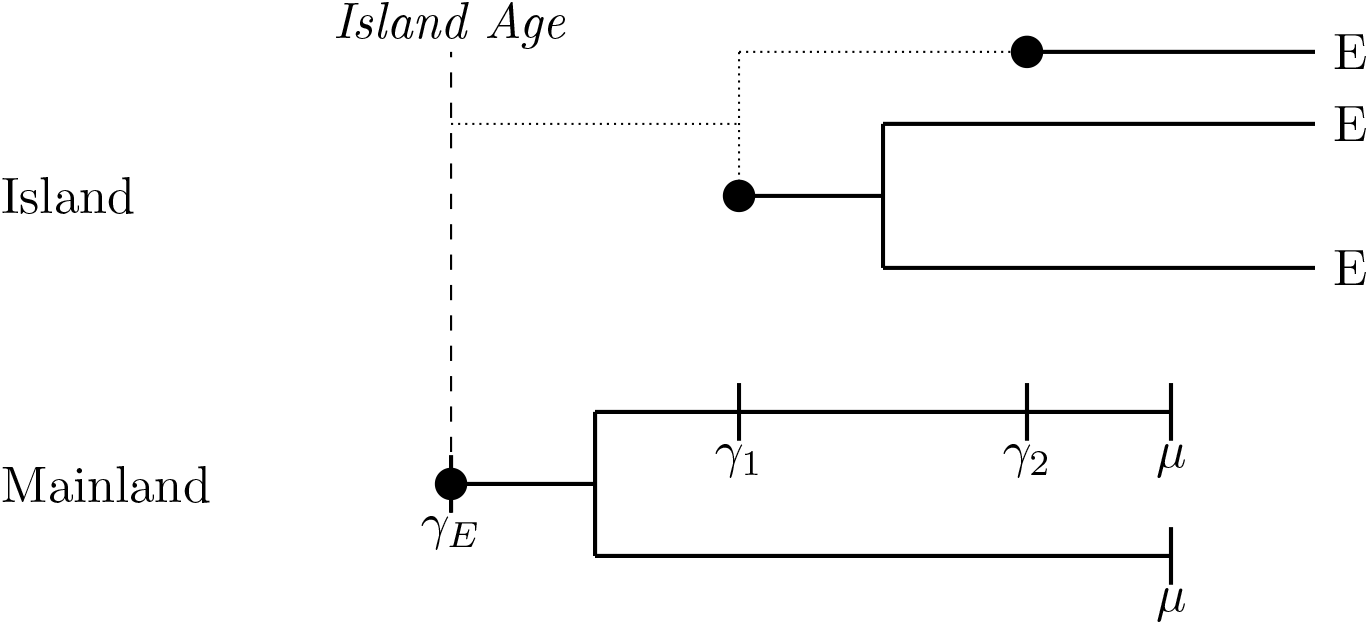
If the mainland species speciates and then one of the descendants colonises the island twice and then both descendants go extinct, and the first island species undergoing cladogenesis before the second immigration event, both colonists are endemic in the *empirical* case they are thought to form a single clade with a colonistation time at maximum island age. *Ideal* data is assigned stac 3 and *empirical* data are assigned stac 6.

In the case of multiple colonisations by the same mainland species, we assume that if the island species has not undergone cladogenesis or anagenesis and is thus still a non-endemic species, the new colonisation will overwrite the colonisation time of the first colonisation. The underlying reasoning is that gene flow from the new colonists will erase the genetic signature that the species was on the island before. If the island species has undergone cladogenesis or anagenesis and is thus an island endemic (or multiple island endemics in a clade), the mainland species can immigrate forming a non-endemic species on the island. In the case of a mainland branching event both descendent species are considered new species and so a re-immigration of the mainland ancestor cannot occur. Therefore, only scenarios where re-immigration without mainland speciation or re-immigration before a mainland speciation are considered. The effect of cladogenesis or anagenesis on re-colonisation is the same so only cladogenesis is shown in the section below, but all scenarios result in the same outcome in cases where anagenesis replaces the island branching time in the scenarios below.

### Colonisation of a different mainland species after cladogenesis on the mainland with no island events

**Figure A36:**
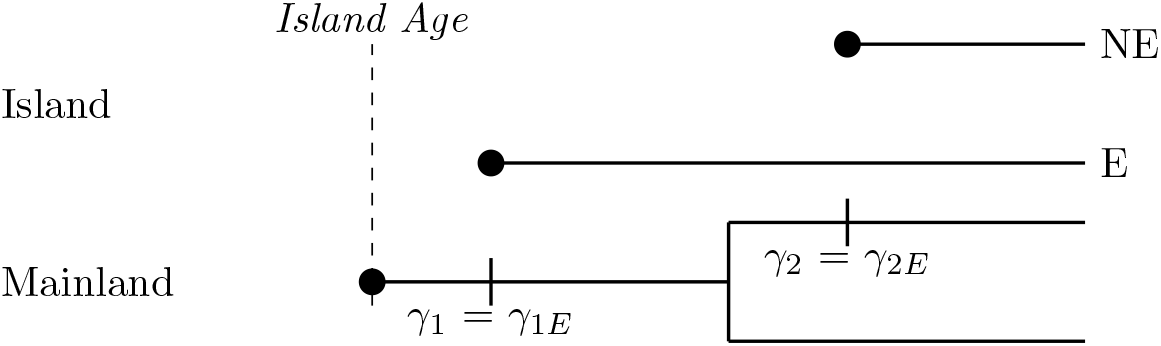
If the mainland species immigrates to the island, then undergoes speciation and one of the descendant species immigrates to the island and both descendants do not go extinct and no events happen on the island, the first island colonist is endemic (E), and the second island colonist is non-endemic (NE). In the *ideal* data the two immigration events are stored as two island clades. In the *empirical* data the first colonisation time (*γ*_1*E*_) is the same as the colonisation time in the *ideal* data (*γ*_1_), and the second colonisation time (*γ*_2*E*_) is the same as the colonisation time in the *ideal* data (*γ*_2_). In the *ideal* and *empirical* data the first colonist is assigned stac 2 and the second colonist is assigned stac 4.

**Figure A37:**
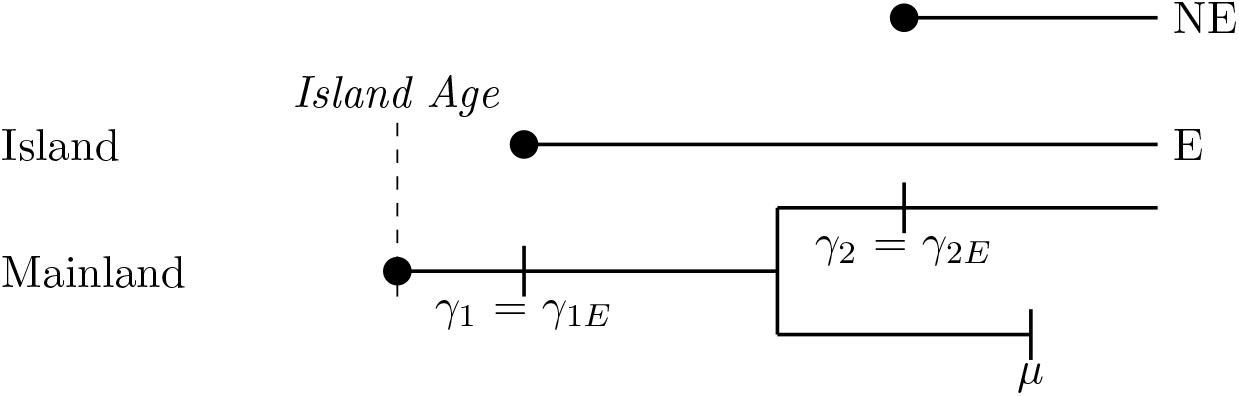
If the mainland species immigrates to the island, then undergoes speciation and one of the descendant species immigrates to the island and survives to the present on the mainland and the other descendant goes extinct and no events happen on the island, the first island colonist is endemic (E), and the second island colonist is non-endemic (NE). In the *ideal* data the two immigration events are stored as two island clades. In the *empirical* data the first colonisation time (*γ*_1*E*_) is the same as the colonisation time in the *ideal* data (*γ*_1_), and the second colonisation time (*γ*_2*E*_) is the same as the colonisation time in the *ideal* data (*γ*_2_). In the *ideal* and *empirical* data the first colonist is stored as stac 2 and the second colonist is stored as stac 4.

**Figure A38:**
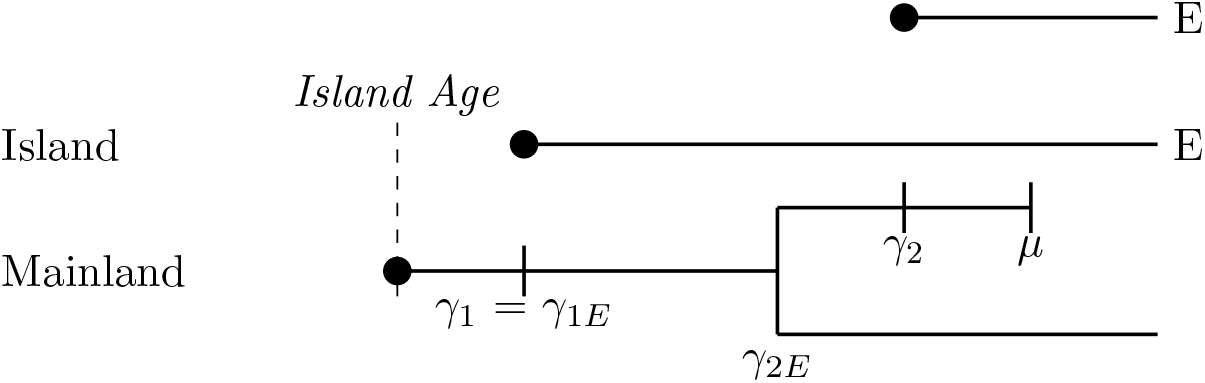
If the mainland species immigrates to the island, then undergoes speciation and one of the descendant species immigrates to the island and goes extinct on the mainland and the other descendant survives to the present, and no events happen on the island, both island colonists are endemic (E). In the *ideal* data the two immigration events are stored as two island clades. In the *empirical* data the first colonisation time (*γ*_1*E*_) is the same as the colonisation time in the *ideal* data (*γ*_1_), and the second colonisation time (*γ*_2*E*_) is the branching time on the mainland. In the *ideal* and *empirical* data each colonist is stored as stac 2.

**Figure A39:**
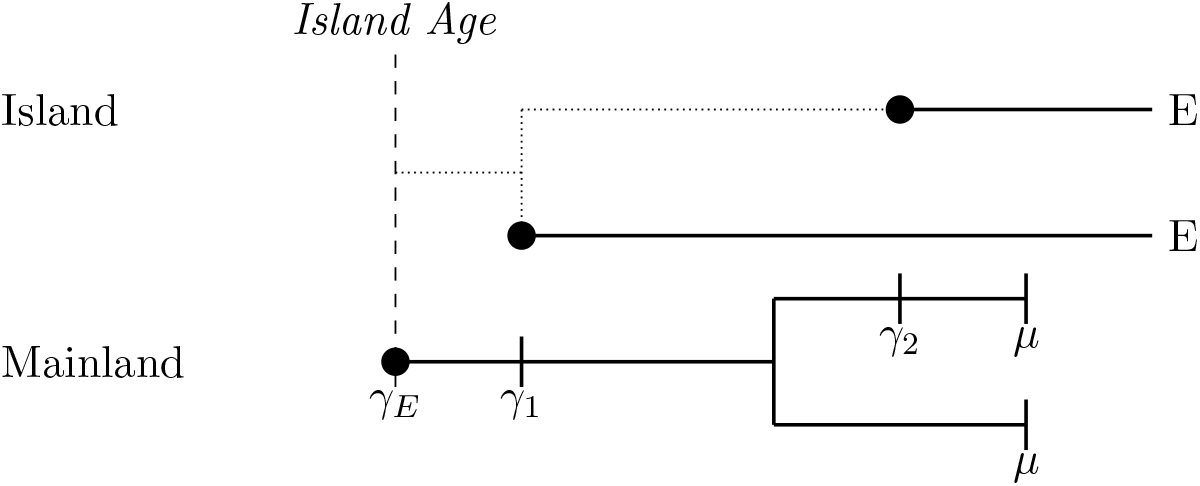
If the mainland species immigrates to the island, then undergoes speciation and one of the descendant species immigrates to the island and both descendants go extinct on the mainland, and no events happen on the island, both island colonists are endemic (E). In the *ideal* data the two immigration events are stored as two island clades. In the *empirical* data they are thought to for, a single clade with a colonisation time (*γ*_*E*_) is the maximum age of the island. In the *ideal* data each colonist is stored as stac 2. In the *empirical* data the island clade is stored as stac 6.

**Figure A40:**
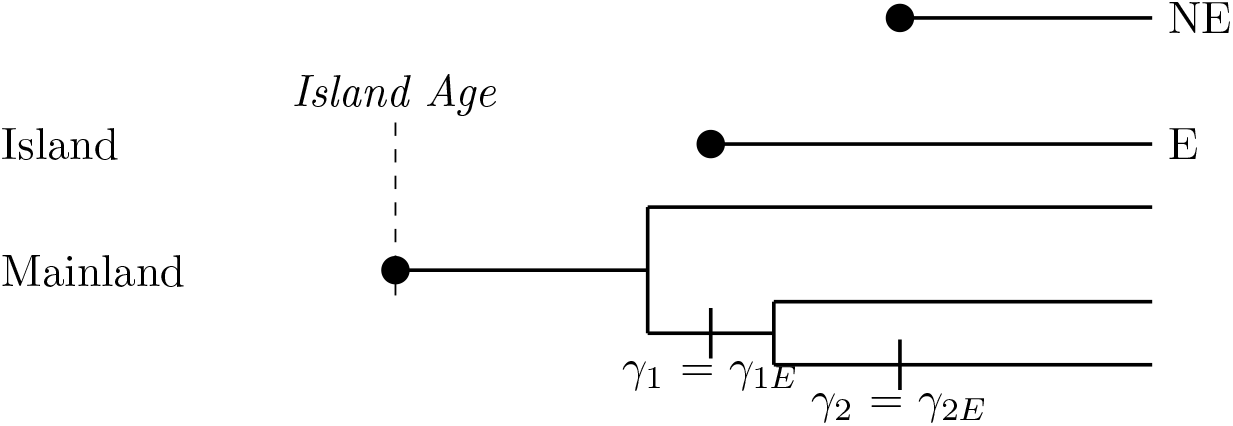
If the mainland species undergoes speciation and one descendant immigrates to the island, then undergoes speciation and one of those descendant species immigrates to the island and both descendants do not go extinct and no events happen on the island, the first island colonist is endemic (E), and the second island colonist is non-endemic (NE). In the *ideal* data the two immigration events are stored as two island clades. In the *empirical* data the first colonisation time (*γ*_1*E*_) is the same as the colonisation time in the *ideal* data (*γ*_1_), and the second colonisation time (*γ*_2*E*_) is the same as the colonisation time in the *ideal* data (*γ*_2_). In the *ideal* and *empirical* data the first colonist is stored as stac 2 and the second colonist is stored as stac 4.

**Figure A41:**
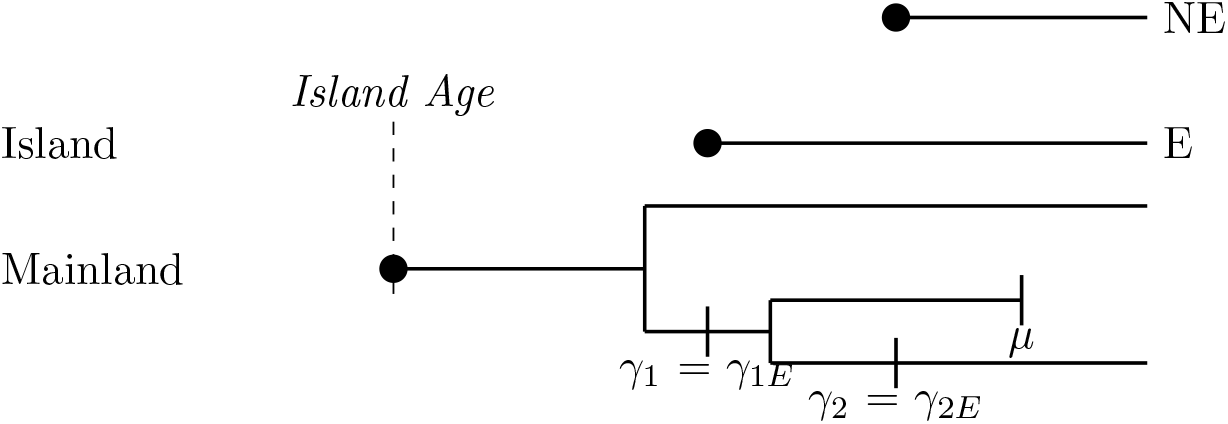
If the mainland species undergoes speciation and one descendant immigrates to the island, then undergoes speciation and one of those descendant species immigrates to the island and the other descendant goes extinct and no events happen on the island, the first island colonist is endemic (E), and the second island colonist is non-endemic (NE). In the *ideal* data the two immigration events are stored as two island clades. In the *empirical* data the first colonisation time (*γ*_1*E*_) is the same as the colonisation time in the *ideal* data (*γ*_1_), and the second colonisation time (*γ*_2*E*_) is the same as the colonisation time in the *ideal* data (*γ*_2_). In the *ideal* and *empirical* data the first colonist is stored as stac 2 and the second colonist is stored as stac 4.

**Figure A42:**
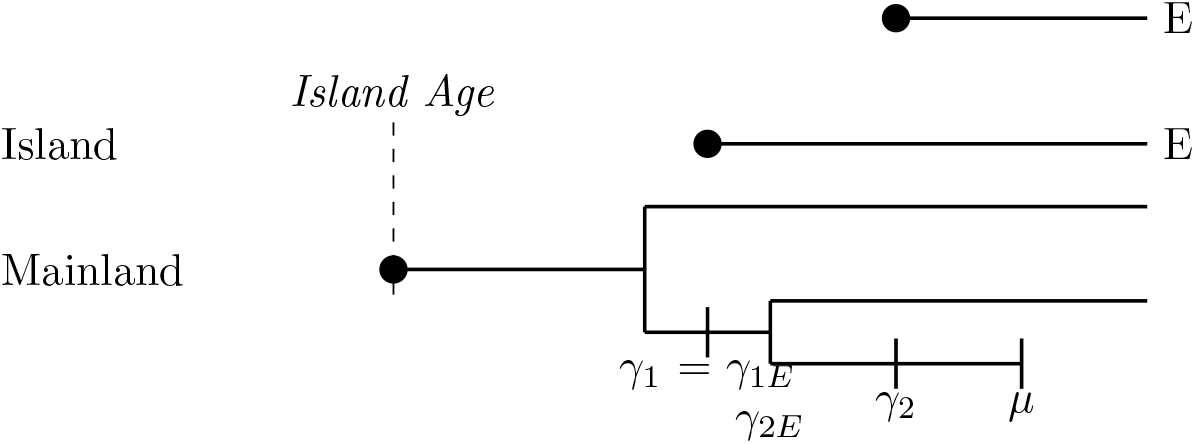
If the mainland species undergoes speciation and one descendant immigrates to the island, then undergoes speciation and one of those descendant species immigrates to the island then goes extinct and the other descendant survives to the present and no events happen on the island, both island colonists are endemic (E). In the *ideal* data the two immigration events are stored as two island clades. In the *empirical* data the first colonisation time (*γ*_1*E*_) is the same as the colonisation time in the *ideal* data (*γ*_1_), and the second colonisation time (*γ*_2*E*_) is the last branching time on the mainland. In the *ideal* and *empirical* data each colonist is stored as stac 2.

**Figure A43:**
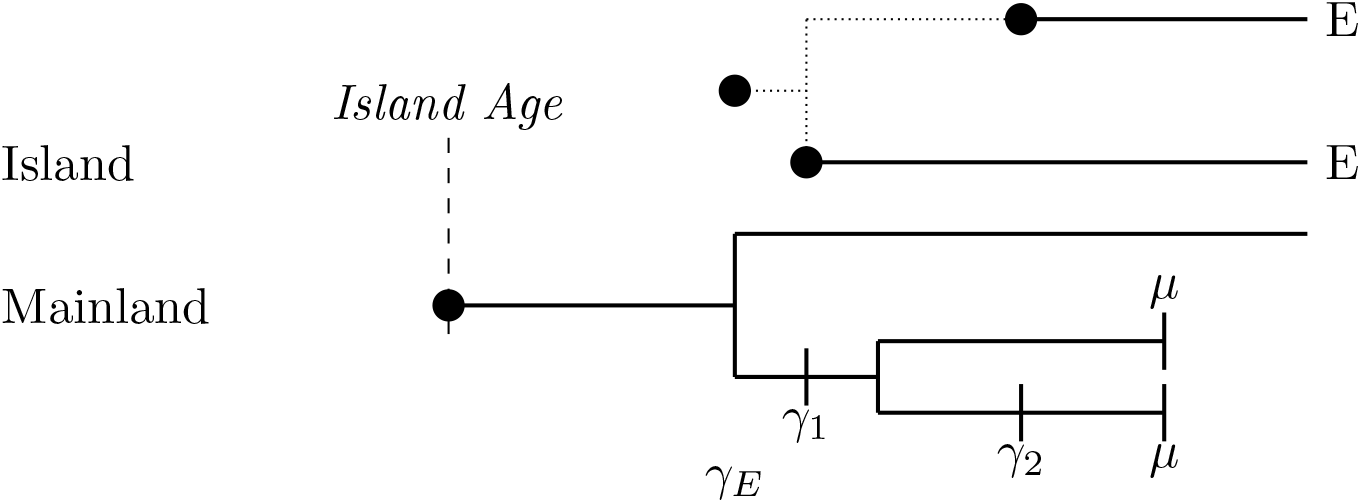
If the mainland species undergoes speciation and one descendant immigrates to the island then undergoes speciation and one of those descendant species immigrates to the island then both descendants go extinct and no events happen on the island, both island colonists are endemic (E). In the *ideal* data the two immigration events are stored as two island clades. In the *empirical* data they form a single clade both colonisation times (*γ*_1*E*_ and *γ*_2*E*_) with a colonisation time at the first branching time on the mainland. In the *ideal* each colonist is stored as stac 2. In the *empirical* data the clade is stored as a stac 2.

Scenarios with more than one colonisation from different species from the same mainland clade with anagenesis or cladogenesis on the island can be determined based on the scenarios outlined above.

## Appendix B

### Stac classification algorithm

The stac is the status of colonist. This is used to inform the DAISIE inference model of the endemicity status of the singleton lineage or the clade, whether it is a *maximum island age colonisation*, or whether there is a recolonisation of the same mainland species. The difference between the *ideal* and *empirical* data sets were explained given numerous different scenarios (note the scenarios above are not exhaustive and only give an overview of the simple cases). We developed an algorithm which can classify the endemicity status and colonisation time of each clade. This section gives a graphical explanation of this algorithm in the form of a decision tree. This algorithm is implemented in the R package DAISIEmainland (Lambert et al., 2021).

**Figure A44:**
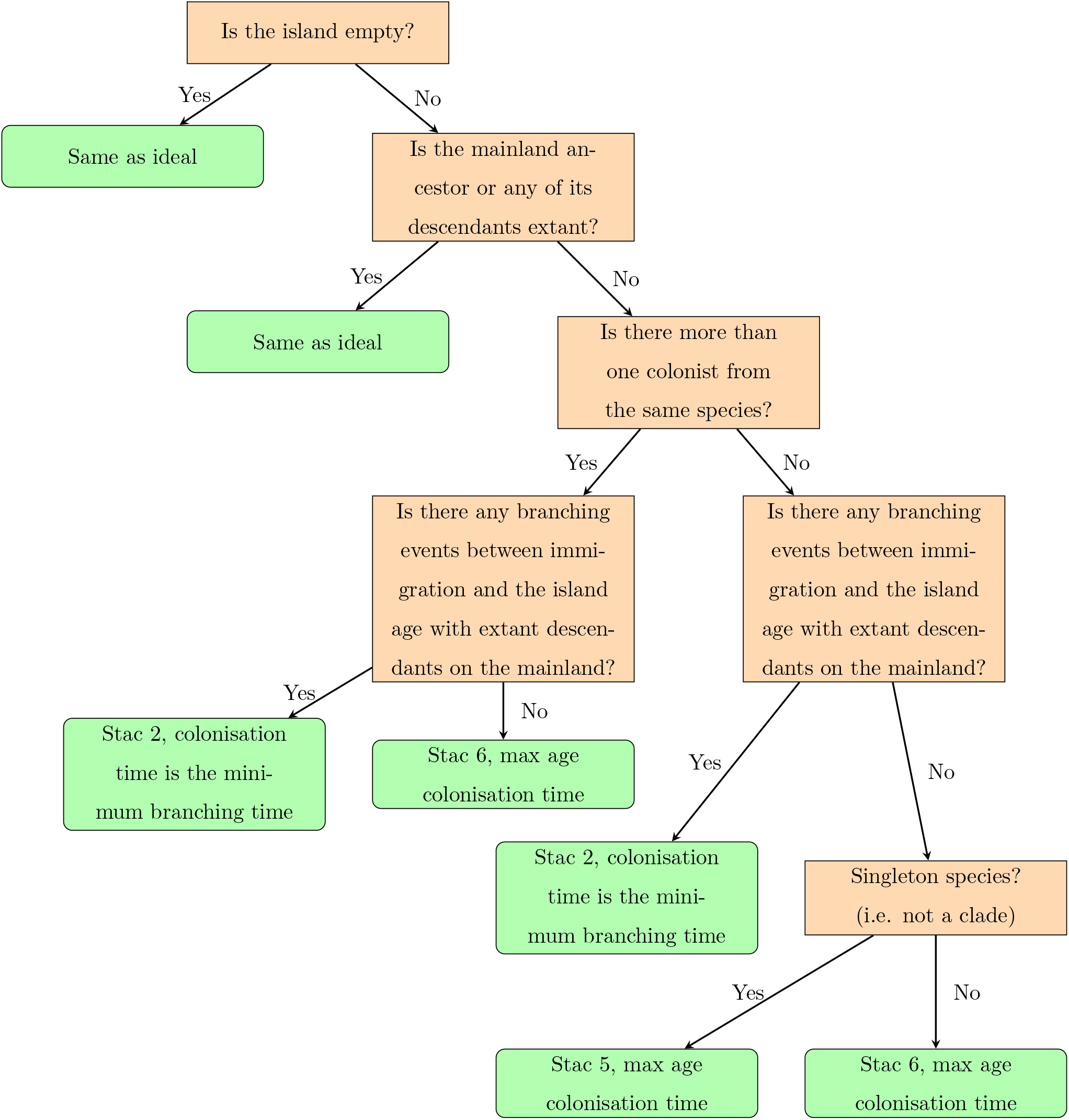
Decision tree showing the classification of the *empirical* data set as compared to the *ideal*. The internal (decision) nodes are shown in orange and terminal (outcome) nodes are shown in green. The outcome of each decision is binary (yes or no).

If the mainland species forms a clade, then different species can immigrate to the island that share the same mainland common ancestor. This is not considered in the DAISIE inference model, given the assumption that mainland species are single independent lineages that cannot undergo speciation or extinction. In these cases, when two or more different species immigrate to the island (as shown below), they evolve under the same diversity-dependent process. However, these are input into the DAISIE inference model as two island clades evolving under independent diversity-dependent processes. The effect of violating this assumption of the DAISIE inference model - along with the change in colonisation time - is one of the focal points of this study, and has empirical importance because mainland species are known to be phylogenetically non-independent, with some clades potentially sharing a recent common ancestor and thus likely to compete under the same diversity-dependent regime on the island. The scenarios below are generalisations of all scenarios of colonisations of different species from the same mainland clade, and can be extrapolated to any case with two or more colonisations of different species from the same clade.

